# Ambivalent roles of nonsense-mediated mRNA decay in heat stress tolerance

**DOI:** 10.64898/2026.10.30.763111

**Authors:** Syed Hussam Abbas, Henrik Mihály Szaker, Éva Darkó, Tünde Erős, Dániel Silhavy, Tibor Csorba

**Affiliations:** Genetics and Biotechnology Institute, Department of Plant Biotechnology, Hungarian University of Agriculture and Life Sciences, 2100, Gödöllő, Hungary; Plant and Horticulture Program, Doctoral School of Agricultural and Food Sciences, Hungarian University of Agriculture and Life Sciences (MATE), 2100, Gödöllő, Hungary; HUN-REN, Agricultural Institute, Centr for Agricultural Research, 2462, Martonvásár, Hungary; HUN-REN, Plant Photo and Chronobiology group, Biological Research Centre, Szeged

**Keywords:** *Nonsense-mediated decay*, *heat stress memory*, *HsfA2*, *HsfA3*, *Arabidopsis*

## Abstract

RNA homeostasis is essential for gene regulation and cellular survival; however, how RNA degradation pathways contribute to stress adaptation remains incompletely understood. Here, we investigated the role of the nonsense-mediated mRNA decay (NMD) in regulating heat stress (HS) memory in a long-term acquired thermotolerance assay. NMD-deficient *Arabidopsis thaliana* mutants *upf1-5* and *upf3-1* exhibited a biphasic phenotype: they displayed transiently elevated heat tolerance during early recovery but became more sensitive to heat during late recovery. Genetic analysis of *upf3;hsfa2* and *upf3;hsfa3* double mutants demonstrated that the early heat tolerance occurs independently of both HsfA2 and HsfA3 HS- memory transcription factors. Under non-treated conditions, the *upf3-1* transcriptome exhibited baseline priming of heat-shock responsive transcripts, likely accounting for the initial heat tolerance. Further transcriptome profiling revealed that *upf3-1* mutants exhibited reduced levels of the functional *HsfA2* isoform (*HsfA2-I*) and attenuated expression of the HsfA2- dependent thermomemory regulon following the triggering stress. NMD deficiency also upregulated immune-response genes while repressing developmental and cell-wall-associated pathways. Together, impaired HsfA2-dependent thermomemory and a compromised growth– defense balance underlies the late heat sensitivity of NMD-deficient plants. Overall, NMD acts as a dual-action post-transcriptional regulator that coordinates the transition from stress response to growth resumption.

**Highlight statement:** Arabidopsis NMD Mutants Exhibit a Biphasic Heat Stress Response, with Early HsfA2/A3-Independent Tolerance and Late HsfA2-Dependent Sensitivity.

## Introduction

High temperature is a major abiotic stress factor that impacts the viability and reproductive fitness of all organisms. Exposure to heat stress (HS) inflicts cellular damage that must be prevented or, if already present, reversed. The collection of pathways dedicated to this mitigation is termed the Heat Shock Response (HSR). HSR operates on many layers, a central level of which is transcriptional regulation. Upon HS a large part of developmental genes is shutdown, while the stress-responsive genes are hyper-transcribed (Akerfelt, Morimoto, & Sistonen, 2010; Charng, Mitra, & Yu, 2023; H. C. Liu & Charng, 2013; Obermeyer et al., 2023; Ohama, Sato, Shinozaki, & Yamaguchi-Shinozaki, 2017; Szadeczky-Kardoss et al., 2022; Yoshida et al., 2011).

The induction of the HSR is a multi-layered process that begins with the activation of Heat Stress Factor A1 family transcription factors (in *Arabidopsis* HsfA1a, A1b, A1d and A1e). Once triggered, HsfA1s launch a transcriptional cascade of secondary HSFs, and other HS- transcripts including heat stress proteins (HSPs), antioxidants, etc. (Akerfelt et al., 2010; Charng et al., 2023; H. C. Liu & Charng, 2013; Ohama et al., 2017). Besides the immediate HSR, plants are capable to remember heat exposures and become primed (acclimated) for a subsequent thermal stress; the process is called thermomemory (or long acquired thermotolerance, LAT). For thermomemory activation the plant utilizes dedicated targets of HsfA1s, the HsfA2 and HsfA3 factors (Charng et al., 2007; Friedrich et al., 2021; Kappel et al., 2023). The HsfA2 and A3 proteins establishes heat memory through promoting the deposition of epigenetic marks that persist on target loci for days and ensure that thermomemory genes remain active (type I memory genes) or are reactivated more robustly (type II memory genes) upon a second heat exposure (Friedrich et al., 2021; Weng et al., 2014). NMD (Nonsense-Mediated mRNA Decay) is a translation-coupled RNA quality control system conserved in eukaryotes, which is essential for development and stress responses; NMD contributes broadly to transcriptome homeostasis, it eliminates aberrant and wild-type transcripts having NMD inducing features (F. Kerenyi, Wawer, Sikorski, Kufel, & Silhavy, 2013; Z. Kerenyi et al., 2008; Kertesz et al., 2006; Kurihara et al., 2009; J. P. Lloyd & Davies, 2013; J. P. B. Lloyd, 2018; Luha, Rana, Vainstein, & Kumar, 2024; Nasim, Karim, Blilou, & Ahn, 2025; Ohtani & Wachter, 2019). The process is activated against mRNAs in which the translational machinery identifies a premature termination codon (PTC). PTC recognition is based on the presence of exon-junction complexes (EJC) located in the 3’ untranslated region (3’UTR), and an unusually long 3’UTR. Besides, if a short open reading frame is present in the 5’UTR (upstream ORF, uORF) before the main protein-coding frame, it also triggers NMD- mediated mRNA decay (Li, Srivastava, Howell, Chudalayandi, & Severin, 2026; Luha et al., 2024). The alternatively spliced (AS) mRNA isoforms are a major source of PTC (Kurihara et al., 2009). The core NMD *trans*-factors are the UP-Frameshift 1 (UPF1), UPF2, UPF3, and the late-acting Suppressor with Morphogenetic effects on Genitalia 1 (SMG1) and SMG7 proteins (F. Kerenyi et al., 2013; Z. Kerenyi et al., 2008; Kertesz et al., 2006; Kurihara et al., 2009; J. P. B. Lloyd, 2018; Ohtani & Wachter, 2019). Plant NMD is autoregulated, the SMG7 mRNA is a direct NMD target (Z. Kerenyi et al., 2008). In *A. thaliana*, a complete loss of function of UPF genes leads to seedling lethality, while hypomorphic *upf* mutants exhibit various developmental abnormalities (Arciga-Reyes, Wootton, Kieffer, & Davies, 2006).

It was shown that NMD regulates several Hsfs, including HSFA2, through the destabilization of AS isoforms. The *HsfA2-I*, the functional form, encoding the HsfA2 protein, is not the target of NMD. The *HSFA2-II*, a non-functional mRNA isoform (Sugio, Dreos, Aparicio, & Maule, 2009), and the *HSFA2-III*, which encodes a short peptide named S-HsfA2, are both targets of NMD (J. Liu et al., 2013). S-HsfA2 feedback regulates positively the HSFA2 locus transcriptional activity (Chen et al., 2025; J. Liu et al., 2013). Based on these, it was implied that NMD fine-tunes plant thermomemory through the HsfA2 pathway, but this hypothesis was not tested experimentally. Neither is known whether NMD regulates other aspects of heat stress memory through HsfA3, or if the impact of AS-NMD on other proposed Hsf targets, including HsfA4c, A7b, B1, B2a (J. Liu et al., 2013)) is biologically relevant.

In the present work, we demonstrate that NMD has an ambivalent role during thermal resilience. On one hand (i) NMD activity establishes a non-stress status at ambient condition through repression of unneeded HS-inducible transcripts, while on the other hand it promotes thermomemory through (ii) increasing *HsfA2-I* and consequently downstream HsfA2- (but not HsfA3-) regulon expression, (iii) by suppressing aberrant induction of the immunity pathway during HS, and (iv) by promoting growth through cell wall and auxin routes during recovery period.

In sum, our work untangles the complex roles of NMD during thermal adaptation and shows that RNA quality control -coupled turnover is a crucial determinant of efficient stress defense and recovery.

## Materials and methods

### Plant materials

*Arabidopsis thaliana* seeds were surface sterilized by bleach treatment, stratified two days at 4°C in the dark and germinated on MS (Duchefa M0222) medium agar plate (1x MS salts, 1% glucose, 1% agar, pH 5.8). Plants were routinely grown in a Sanyo MLR-350 growth chamber under cool white light at 21°C long-day condition photoperiod.

### Mutant lines used in the study

T-DNA insertion lines *upf1-5* (SALK_081178), *upf3-1* (SALK_025175), *hsfa2-1* (SALK_008978) *and hsfa3-1* (SALK_011107) were obtained from NASC (https://arabidopsis.info). Double mutants were generated by crossing the respective single mutants, and homozygous lines were confirmed by genotyping.

### Genotyping

Genomic DNA was extracted using a nucleic acid extraction buffer (100 mM glycine, 10 mM EDTA, 100 mM NaCl, 2% SDS), purified with phenol-chloroform-isoamyl alcohol (25:24:1, pH 8.0), precipitated in 96% ethanol and resuspended in sterile water. Genotyping PCR was performed using Taq DNA polymerase (NEB, M0273S) according to the manufacturer’s instructions. Genotyping primers were described previously (Friedrich et al., 2021; Szadeczky- Kardoss et al., 2022; Szaker et al., 2019; Szaker et al., 2025).

### Heat stress treatments

For the long acquired thermotolerance (LAT) heat stress treatment, the 4 days-old seedlings were primed with two-step priming (P) protocol in a water bath in the dark: 37°C/60 min (ZT6- ZT7), 21°C/90 min, and at 44°C/45 min (ZT8,5-ZT9,5). Three days after acclimation, primed seedlings were exposed to a triggering (P/T) heat treatment at 44°C/50-80 min in the water bath. Seedlings were returned to 21°C for post-stress recovery (P/T+recovery). Experiments were performed in at least three independent replicates. For molecular analyses, seedlings were harvested immediately after the priming (P) step and following one day of recovery (P+1d) at 21°C (RT-qPCR assay), or after priming and treatment (P/T) and P/T followed one day (P/T+1d) recovery (RNA-seq). Samples were flash-frozen in liquid nitrogen and stored at −80°C. For physiological characterization, and for phenotyping and statistical analysis, plants on agar plates were analyzed 4 days (P/T+4d) or 14 days (P/T+14d) after the priming and triggering treatment (P/T), and in parallel non-treated controls (NT, plant age was 10, 12 or 20 days old, respectively) as shown in figure 1.

**Figure 1:**
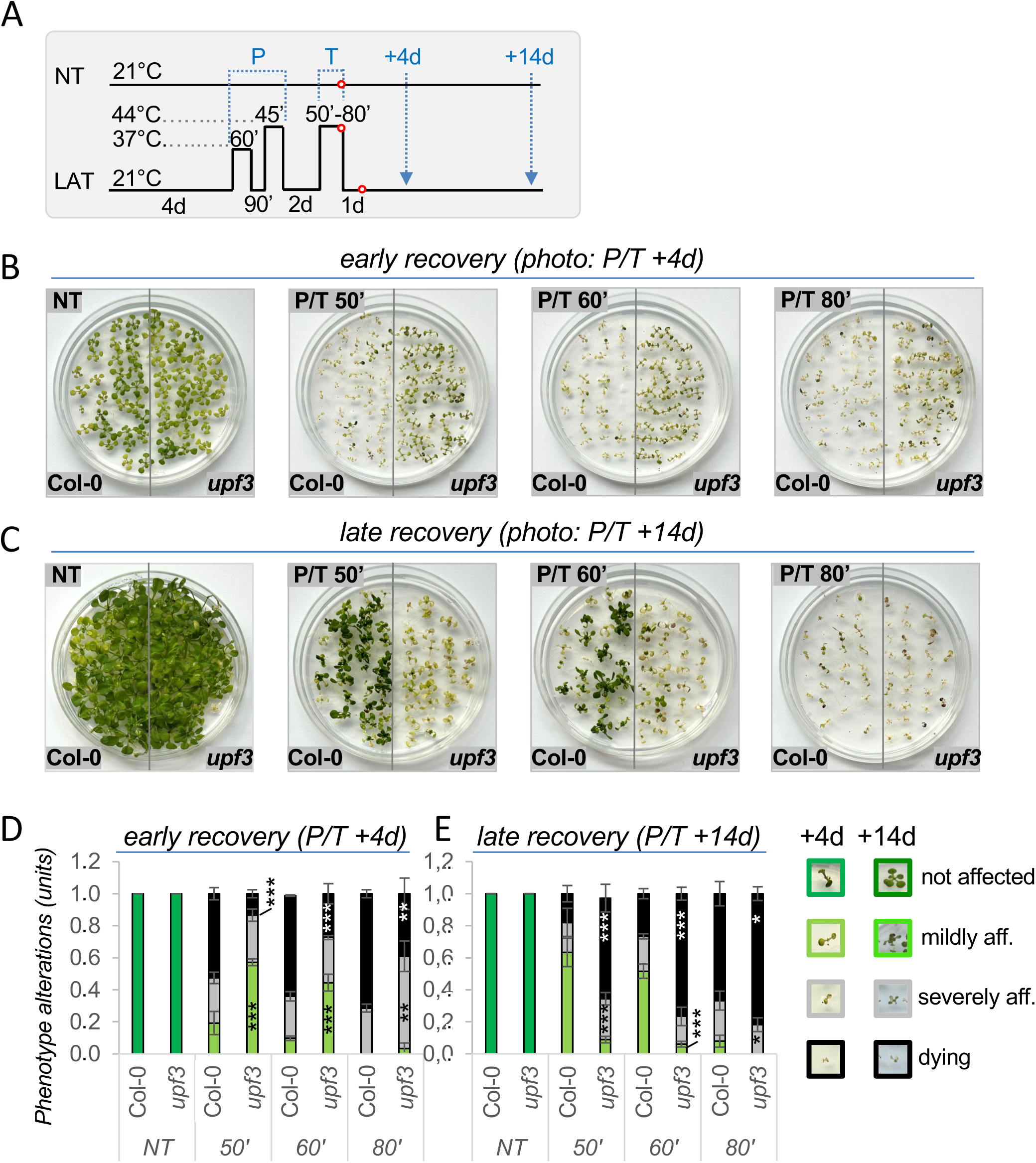
NMD has ambivalent roles during heat stress response. (A) Treatment scheme for long- acquired thermotolerance (LAT) regime to test HS tolerance; priming (P: 37°C/60′, 21°C/90′, 44°C/45′) was followed by 2 days interval at 21°C, than triggering (T) heat treatment for 50′, 60′, or 80′ (P/T); red cicles show sample collection times for transcriptome analysis (*see latter*); (B) Phenotypes of wild-type (Col-0) and mutant *upf3* taken on day 4 post-treatment (P/T+4d, early recovery) or (C) 14 days post-treatment (P/T+14d, late recovery). (D) Quantification of early phenotypes (+4d) shown in *panel B*; (E) quantification of late phenotypes (+14d) shown in *panel C*; seedlings were scored as not affected, mildly affected, severely affected, or dying (*see legend on right for each*). Data are mean ± SD of three independent experiments, and asterisks indicate significant differences to Col-0 (*p < 0.05; **p < 0.01; ***p < 0.001, unpaired two-sided t-test).

### Phenotypic scoring

Seedling phenotypes were scored visually at early (+4d) and late (+14d) recovery after P/T, alongside non-treated controls grown on the same plate. Each seedling was assigned one of the four categories: not affected (green, expanded cotyledons and true leaves), mildly affected (partial wilting, reduced leaf expansion or brownish-green pigmentation), severely affected (extensive wilting and chlorosis, with some green tissue remaining) or dying (withered, yellow- brown cotyledons and true leaves without green tissue). For each genotype and treatment, an average of 30 seedlings per plate were scored, and experiments were repeated at least three times independently. The proportion of seedlings in each category was calculated per plate, and differences between genotypes were tested using an unpaired two-tailed Student’s t-test (P < 0.05).

### RNA extraction and RT-qPCR

Seedlings (about 30 mg) were flash-frozen in liquid nitrogen and stored at −80°C until the next step of the process. Total RNA was extracted from the frozen seedling tissue ground to a fine powder in liquid nitrogen using a Quick-RNA Miniprep kit with on-column DNase I digestion (ZYMO Research, Irvine, CA). First-strand cDNA was synthesized from 1 μg total RNA using random primers and the ProtoScript II First Strand cDNA Synthesis Kit (NEB, E6300S, https://www.neb.com) according to the manufacturer’s protocol. Quantitative PCR (qPCR) was performed using qPCR Master Mix (NEB, M3003S, https://www.neb.com) and qPCR reactions were run in LightCycler96 real-time PCR instrument (Roche). At least three independent biological replicates were generated per genotype and treatment condition; error bars represent the standard error of the mean. *P*-values were calculated using unpaired two- tailed Student’s *t*-test to assess the significance of differences. Differences were considered statistically significant at *P* < 0.05. All comparisons were made against Col-0 within the same treatment condition unless otherwise stated. Gene expression levels were normalized to the reference gene *PP2AA3 (AT1G13320)*, previously extensively proved as a very stable transcript under both control ambient or heat stress conditions (Karanyi et al., 2022; Szadeczky- Kardoss et al., 2022; Szaker et al., 2025). Genotyping primers were described previously (Friedrich et al., 2021; J. Liu et al., 2013; Sugio et al., 2009; Szadeczky-Kardoss et al., 2022; Szaker et al., 2019; Szaker et al., 2025).

### Physiological measurements

Chlorophyll *a* fluorescence measurements were performed on intact plants using IMAGING PAM Maxi (Walz, Effeltrich, Germany), as described previously (Darko *et al*., 2011), except that 56 umol m^-2^ s^-1^ actinic light intensity was used for 15 min to induce photosynthesis (Lichtenthaler, 1987) after 20 min dark adaptation period. The fluorescence parameters were determined according to (Lichtenthaler, 1987). Chlorophyll (a+b) content was determined using a Cary-100 UV-VIS spectrophotometer (Varian, Middelburg, Netherlands) after extraction of 50 mg shoot plantlets in 80% acetone, using the method of (Lichtethaler, 1987). Electrolyte leakage (EL%) was determined as described previously by (Asghar *et al*., 2021) with the use of 0.05 shoot plants for each sample. Six biological repetitions were used for each genotype and treatment.

### Transcriptome analysis

Col-0 (wild-type) and *upf3-1* mutant plants were profiled by RNA-seq under three conditions: non-treated (NT), heat priming+triggering (P/T), and one day of recovery after triggering (P/T +1d). Raw RNA-seq reads were trimmed using Cutadapt (v4.7.) (Martin, 2011). The trimmed reads were aligned to the *Arabidopsis thaliana* TAIR10 (v62.) reference using HISAT2 and were processed with SAMtools (v1.22.1.) to generate BAM files (Danecek et al., 2021). Gene- level raw read counts were assembled into a count matrix with featureCounts (v2.0.8.) (Liao, Smyth, & Shi, 2014) for downstream differential expression analysis with DESeq2 (v1.50.2.) (Love, Huber, & Anders, 2014). Differential expression between *upf3-1* and wild-type Col-0 was assessed within each condition (log_2_FC > 1, BH adjusted p < 0.05).

Gene Ontology (GO) term enrichment was performed with clusterProfiler (v4.18.4.) (T. Wu et al., 2021). GO terms with a BH adjusted p-value below 0.05 were considered significant. To reduce redundancy among related GO terms, semantic similarity reduction was performed using the Wang similarity measure with a cutoff of 0.7.

For each Araport11 transcript, we computed 3′UTR length, whether an intron overlaps the 3′UTR, and whether an upstream open reading frame (uORF) is present. Each feature was reduced to one value per gene: the longest 3′UTR across isoforms, and an intron or uORF flag if any isoform had one. Experimental uORF evidence came from the RiboTaper (primary analysis) and CiPS (sensitivity analyses) datasets (H. L. Wu et al., 2024). A gene with too little ribosome-profiling signal can’t show that it lacks a uORF. We therefore called a gene Ribo- seq-detectable if its TPM was at least 1 in GSE183264. Genes with a positive uORF call were always kept, and genes without one counted as negatives only if they were detectable. 3′UTR length was compared between expression-status groups with Wilcoxon rank-sum tests. The prevalence of 3′UTR introns and uORFs was compared with Fisher’s exact tests. P-values were adjusted with the Benjamini–Hochberg procedure within each feature family.

Expression trajectories were constructed using DESeq2-normalized counts. Counts were transformed as log_2_(normalized count + 1), and mean expression was calculated for each gene within each of the six genotype–treatment combinations. The 2,000 most variable genes were selected for the primary analysis and each selected gene’s six-condition profile was standardized to a z-score. K-means clustering was performed on these standardized profiles with six clusters, 50 random starts and a maximum of 100 iterations, using a fixed random seed. Cluster trajectories were summarized by genotype and treatment.

For *cis*-regulatory element enrichment, candidate transcription-factor binding motifs were searched in the 1-kb region upstream of gene’s transcription start site; motif presence was compared against the genome-wide promoter background by Fisher’s exact test, reported as odds ratio and significance.

Heat shock response (HSR) gene sets were defined using the treatment-associated DESeq2 contrasts in Col-0. Genes induced under P/T but not P/T+1d were classified as acute or transient responders; genes induced under both P/T and P/T+1d were classified as sustained responders; and genes induced under P/T+1d but not P/T were classified as delayed responders. The broader HSR-responsive set comprised genes induced under either treatment condition, while genes expressed in at least one Col-0 sample but not meeting the induction criteria were classified as other expressed genes. Paired Wilcoxon signed-rank tests were applied to per- gene expression means for descriptive Col-0 versus *upf3* comparisons.

For each HSR gene, Pearson’s correlation coefficient was calculated between its Col-0 wild type expression profile across NT, P/T, and P/T+1d and the corresponding wild type expression profile of HSFA2 or HSFA3. Genes with a Pearson correlation coefficient (*r*) > 0.7 with HSFA2 were assigned to the HSFA2-correlated group, while those with *r* > 0.7 with HSFA3 were assigned to the HSFA3-correlated group. Genes exceeding the threshold for both transcription factors were included in both groups and genes with correlations ≤ 0.7 for both factors were classified as ‘neither-correlated’. HSFA2, the adjacent gene AT2G26140 and HSFA3 were excluded from the target-gene correlation analysis to avoid self-correlation or inclusion of the specified neighbouring locus. The corresponding standardized wild type HSFA2 or HSFA3 expression profile was used as the reference trajectory in the relevant transcription-factor-correlated groups.

A priming index was calculated for HSR-classified genes using the genotype-associated expression effect under NT relative to the largest Col-0 treatment induction estimate. The index was defined as the log_2_FC for *upf3* versus Col-0 under NT divided by the maximum of the Col- 0 log_2_FC fold changes for P/T versus NT and P/T+1d versus NT. Genes were retained for this analysis only when the denominator was finite and positive and the resulting index was finite.

Analyses and figures were performed and generated in R using DESeq2 and the associated Bioconductor and CRAN packages (including clusterProfiler, org.At.tair.db, rtracklayer, readxl, rrvgo, ggplot2, dplyr, tidyr, tibble and patchwork). Unless otherwise specified, differential-expression significance was defined using a Benjamini–Hochberg-adjusted p- value threshold of 0.05. All R scripts used for analysis can be found on https://github.com/TCsorbaGroup/Heat_Stress_Recovery_UPF3_RNAseq_analysis. Raw RNA-seq data have been made available in the SRA repository under the BioProject ID: PRJNA1552212.

## Results

### NMD restricts early heat tolerance but supports long-term recovery

LAT assays were used to evaluate the role of NMD in regulating thermomemory. We subjected the NMD mutants *upf3-1* and *upf1-5* (referred to as *upf3* and *upf1*) to a triggering heat stress assay (T) following a priming (P) treatment (primed and triggered, P/T), and then we monitored plant recovery over a two-week post-stress period (Fig 1A-E and Fig S1). Non-treated (NT) plants were used as controls.

During early recovery (P/T+4d), both NMD mutants displayed a mild heat-tolerant phenotype compared to the wild-type Arabidopsis (Columbia-0, Col-0): their leaves were larger and exhibited less shrinking, though they developed a distinct brownish-green pigmentation (Fig 1B,D and S1A). In contrast, Col-0 plants were more affected at this stage, displaying withered, yellow-brown cotyledons and true leaves.

Notably, the phenotypic difference gradually reversed during the two weeks recovery period, the mutant plants lost their pigmentation and developed a HS sensitivity phenotype (Fig 1C,E). At the end of the two-week recovery, the Col-0 plants recovered much more successfully than the NMD mutants (Fig 1C,E and S1C,D).

To analyze the impact of NMD absence at the physiological level, we performed a suite of measurements at early (P/T+4d) and late stage of recovery (P/T+14d), in accordance with the visual inspection timepoints on Col-0 and *upf3* (being a stronger allele (Arciga-Reyes et al., 2006))(Fig 2). Under ambient conditions, the total chlorophyll content of the *upf3* was slightly lower than that of Col-0 (Fig 2A), but following 4-day recovery the chlorophyll levels decreased more strongly in Col-0 than in *upf3* (Fig. 2A, *left*). During the late recovery (P/T+14d) however, Col-0 plants recovered significantly better (Fig. 2A, *right*).

**Figure 2:**
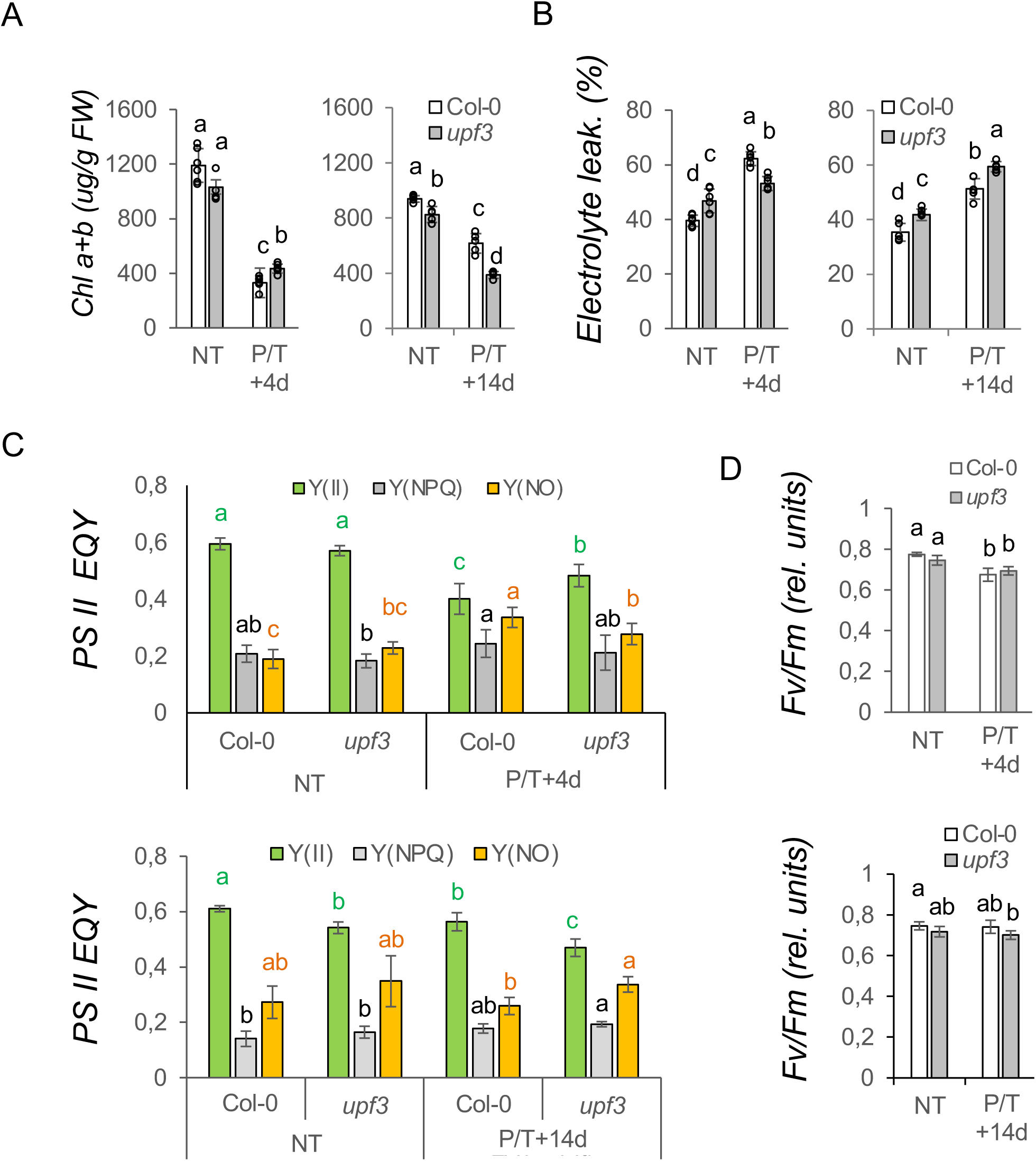
Physiological parameter changes of wild-type (Col-0) and NMD mutant *upf3* plants in response to heat. Treatment was done as shown in Figure 1A. (A) Total chlorophyll content (Chl a+b, µg/g FW); (B) Electrolyte leakage (%); (C) Effective quantum yield of PSII (Y(II)), regulated non- photochemical (Y(NPQ)), and non-regulated non-photochemical (Y(NO)) chlorophyll *a* fluorescence parameters; (D) Optimal quantum yield of PSII (Fv/Fm). All measurements were taken at ambient (non- treated, NT) and following 4 or 14 days of P/T recovery (P/T+4d or P/T+14d, respectively); for details see main text. Data are mean ± SD of six independent experiments (*A, B panels*) or 24 measurements (*C, D panels*), and letters indicate significantly different groups (p<0.05, Tukey’s post-hoc test).

To assess membrane integrity, we measured electrolyte leakage. Under NT conditions *upf3* displayed slightly higher leakage rates (Fig 2B). During recovery, ion leakage increased in both genotypes, but *upf3* exhibited significantly lower leakage at the early recovery stage and higher leakage rates at the late recovery stage (Fig. 2B).

To evaluate photosynthetic performance, we determined key chlorophyll fluorescence parameters. While the effective quantum yield of photosystem II (Y(II)) expectedly declined in response to heat during the early (P/T+4d) recovery phase, the *upf3* mutant retained a significantly higher Y(II) (Fig 2C, *top*). During the later phase of recovery (P/T+14d), the Col- 0 plants recovered more efficiently, as shown by higher Y(II) (Fig.2C, *bottom*). The lower level of Y(II) in *upf3* was accompanied by higher levels of regulated Y(NPQ) and non-regulated Y(NO) non-photochemical quenching compounds (Fig. 2C). The elevated Y(NO) indicates an inability to safely dissipate excess excitation energy, leading to photoinhibition. The maximum photochemical efficiency of PSII (Fv/Fm) was mildly and equally reduced in both genotypes at early recovery and slightly more reduced in the mutant at late recovery stage (Fig. 2D).

Taken together, the visual observations and physiological measurements consistently show that although the NMD mutants are more thermotolerant in the early recovery phase, over a two- week period, the wild-type plants recover much better. Thus, NMD plays a dual role in thermotolerance; it inhibits early but promotes late HS recovery. These data also suggest that at least two contrasting layers of recovery pathways act during the stress-to-regrowth physiological switch.

### NMD remains active during and after heat stress

To compare the molecular changes in wild-type and *upf3* NMD mutant plants during the execution of heat stress memory and very early recovery, we profiled the transcriptomes of non-treated (NT), primed-and-triggered (P/T) and recovering (P/T+1d) plants (Fig. 1A).

First, we studied whether NMD functions during and after the HS on selected NMD *trans* factors. As expected, the UPF3 mRNA showed strongly reduced expression in the *upf3* mutant. The SMG7 mRNA, which is a strong direct target of NMD, was overexpressed in *upf3* under all conditions. By contrast, the UPF1 mRNA was elevated only after HS, and UPF2 was unchanged (Fig 3A). To assess genome wide functions of NMD pathway, we compared the expression of a previously defined core NMD-target set (300 genes, the common targets of *upf1* and *upf3* mutants (Szaker et al., 2025)) Fig. 3B). The *upf3*/Col-0 ratio for NMD targets remained significantly elevated in all states (1.93-, 1.87- and 1.81-fold at NT, P/T and P/T+1d; p < 0.001) indicating that NMD is active throughout the HS-recovery experiment.

**Figure 3:**
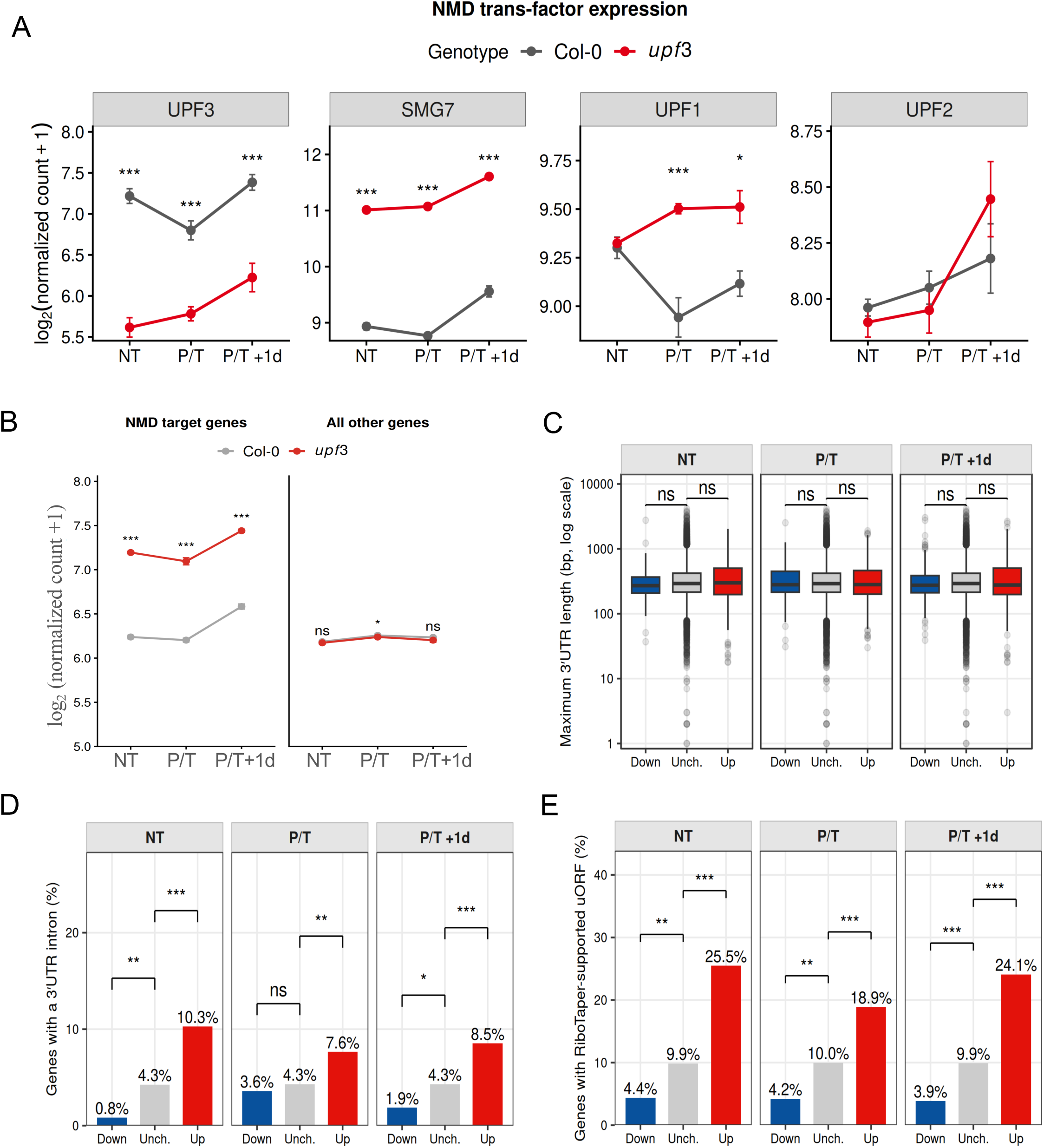
NMD regulation of target genes at ambient (NT), heat stress (P+T) and recovery (P/T+1d). (A) Expression of core NMD *trans* factors (UPF3, SMG7, UPF1, UPF2) in Col-0 (grey) vs *upf3* (red); asterisks indicate genotype significance at each condition, see below. (B) NMD-target derepression: estimated mean expression of previously defined NMD-targets (Szaker et al., 2025) versus non-target genes in Col-0 (grey) and *upf3* (red); asterisks indicate genotype significant difference (Col-0 vs. *upf3*) at each condition (mixed-effects model). (C-E) NMD *cis*-feature analysis of genes differentially down-regulated (Dw; NT n=137, P+T n=313, REC n=265), all expressed genes (All; NT n=23,279, P/T n=22,980, REC n=22,981), and up-regulated (Up; NT n=352, P/T n=475, REC n=552) in *upf3* vs. Col-0 wild-type : (A) 3’UTR length, (B) percentage of genes carrying with intron in their 3’UTR, and (C) percentage of genes carrying an upstream open reading frame (uORF). significance * p<0.05, ** p<0.01, *** p<0.001, ns = not significant, comparing within each condition; Wilcoxon rank-sum test for 3’UTR length, Fisher’s exact test for intron/uORF relative to background.

Next, we analyzed the genes that showed differential expression (DEGs, defined as log_2_FC > 1, adjusted p < 0.05;) in *upf3* plants relative to the wild-type. In all three states, more upregulated than downregulated DEGs were found in *upf3* (n = 606, 547 and 608 upregulated vs. n = 248, 317 and 349 downregulated in NT, P/T and P/T+1d, respectively; Fig. S2A). This is consistent with the role of NMD in destabilizing target transcripts (Gloggnitzer et al., 2014; Riehs-Kearnan, Gloggnitzer, Dekrout, Jonak, & Riha, 2012; Szaker et al., 2025). A significant part of these upregulated transcripts were genotype-dependent (n = 163 up in all conditions), in contrast to the downregulated ones (n = 12) (Fig S2B,C).

The nature of the NMD mechanism suggests that upregulated gene set contains both direct and indirect NMD targets, whereas downregulated genes are always indirect targets. To assess the ratio of direct/indirect NMD targets in the upregulated DEGs (Up), we analyzed the canonical NMD *cis*-features (3′UTR length, intron in the 3′UTR (Z. Kerenyi et al., 2008) or uORF in the 5’UTR (H. L. Wu et al., 2024)) in the Up and Down genes and compared them with the unchanged (Unch.) control genes (Fig. 3C–E). The NMD features were not overrepresented in the Down set (relative to the control), which is consistent with these being indirect NMD targets. By contrast, Up genes had an increasing trend in 3’UTR lengths (NT and P/T+1d, Fig S3C), and were significantly enriched for 3′UTR introns (2.5-, 1.8- and 2-fold at NT, P/T and P/T+1d, vs unchanged set; Fig. 3D) and uORF (2.6-, 1.9- and 2.4-fold at NT, P/T and P/T+1d, vs unchanged set; Fig. 3E) NMD features. The enrichment of NMD features in *upf3* upregulated genes reflects the stabilization of direct targets in the absence of NMD. However, these features were present in only a fraction of upregulated genes (<10.3% 3’UTR intron- and <25.5% uORF-containing genes), indicating that most upregulated genes are not direct NMD targets.

Together, these data show that NMD remains active throughout and after the heat stress regime, and that although its loss stabilizes several direct targets, many of the upregulated transcripts are probably secondarily changed.

### NMD deficiency causes heat-dependent de-repression of immunity pathways

To unravel the molecular basis of the altered heat tolerance, the *upf3* DEGs were analyzed for enriched GO terms (Fig. S2D). Notably, GO studies revealed state-specific responses in *upf3*. In NT plants, the only term enriched among upregulated genes was “RNA modification”, which remained overrepresented at P/T and P/T+1d, suggesting a basal alteration of RNA homeostasis in the absence of NMD. Downregulated genes at NT were moderately enriched for GO terms related to cell wall metabolism and oxidative-stress. After heat (P/T) and during recovery (P/T+1d), upregulated genes in *upf3* were strongly enriched for immunity-related terms (*see detailed list*, Fig S2D). Surprisingly, although *upf3* showed early heat tolerance (Fig1-2), typical heat-response GO terms were not enriched, suggesting that immunity, rather than a canonical HSR, is activated during HS in the NMD-deficient mutant. The downregulated genes at P/T and P/T+1d were enriched for growth- and cell wall-associated terms, predicting a decreased recovery capacity in *upf3* following HS.

Together, these changes are consistent with a shift in the growth–immunity balance toward immunity in the absence of NMD during HS and early recovery.

### Heat-dependent immune activation in *upf3* is consistent with transcriptional induction

Since we did not find HSR differences when *upf3* DEGs were analyzed, we employed a different approach. We partitioned the 2,000 most variable genes of the transcriptome into six clusters (see *Materials and Methods*, Fig 4A) and then these clusters were analyzed. Cluster 1 was enriched for growth- and metabolism-related GO terms, cluster 2 for heat stress and proteostasis terms, cluster 3 for “response to wounding”, and cluster 4 for “RNA modification” and “macromolecule modification” terms. Cluster 5 showed strong enrichment for immunity- related GO terms, whereas cluster 6 did not show any GO enrichment (Fig S3). Importantly, apart from cluster 5, none of the clusters showed strong genotype-specific differences between the wild-type and *upf3* (Fig 4A), indicating that most of the variance of these genes is driven by heat rather than by genotype. Cluster 5, the only group in which gene expression was strongly NMD-dependent, was further analyzed.

**Figure 4:**
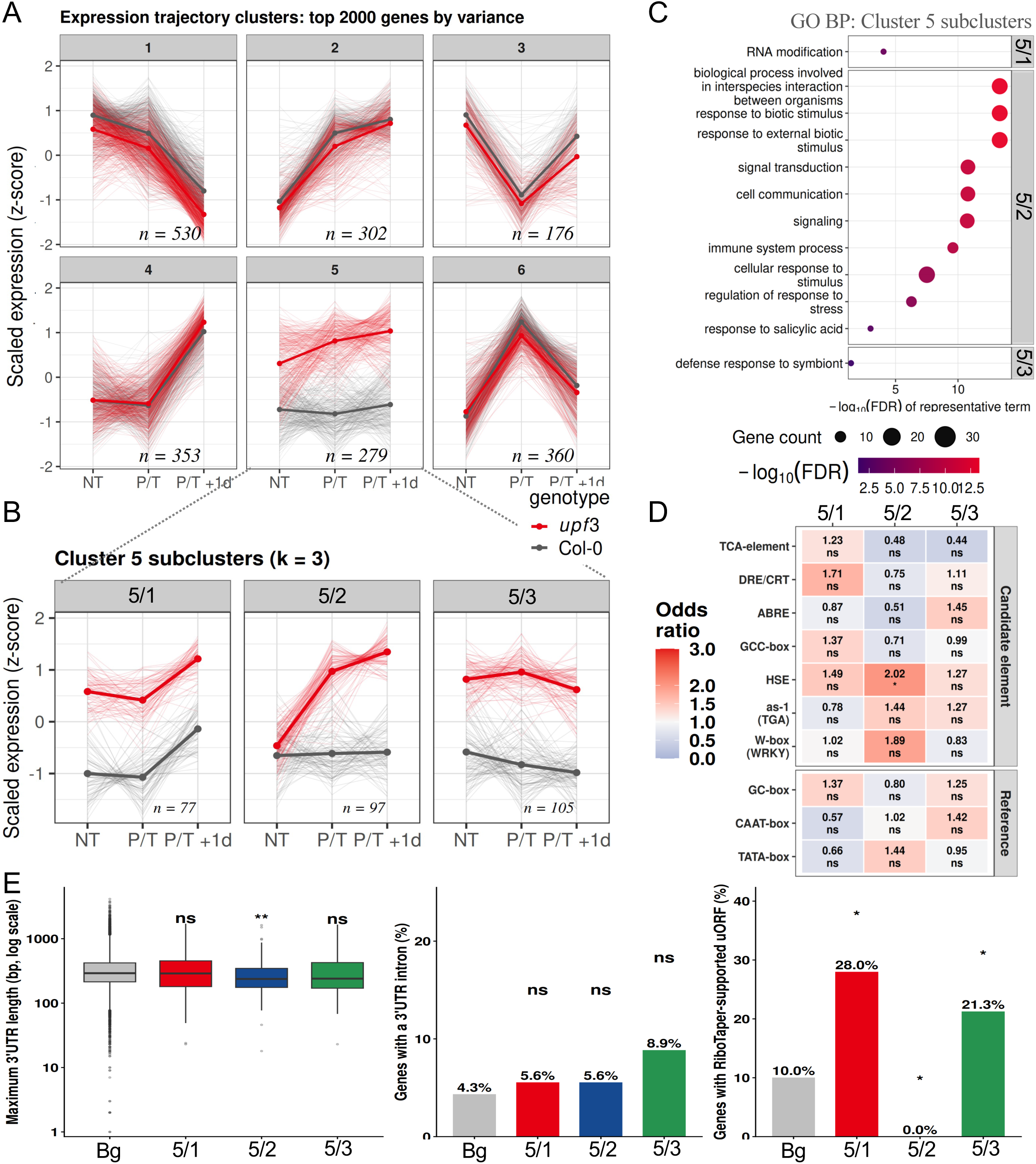
Immunity transcripts accumulate during heat stress in *upf3*. (A) K-means clustering (k=6) of expression trajectories for top 2000 genes-by variance in *upf3* vs. Col-0 plants, based on z-score normalized log_2_(FPKM+1) values under non-treated (NT), heat-stress (P/T), and recovery (P/T+1d); thin lines are individual genes, bold lines show mean values. (B) Sub-clustering of Cluster 5 (k=3); (C) GO enrichment analysis of Cluster 5 subclusters, computed with clusterProfiler (hypergeometric test, BH-adjp<0.05); non-redundant BP terms were obtained by semantic-similarity reduction (rrvgo, threshold 0.7); (D) *Cis*-regulatory element enrichment of 1 kb promoters (relative to TSS) for Cluster 5 subclusters vs. genome background, tested by Fisher’s exact test. Candidate elements: W-box (WRKY-binding), as-1 (TGA-binding, SA-responsive), HSE (heat-shock element), GCC-box (ERF-binding, ethylene-responsive), ABRE (bZIP-binding, ABA-responsive), DRE/CRT (drought/cold-responsive), TCA-element (SA-responsive); core elements used as controls: TATA-box, CAAT-box, GC-box. significance * p<0.05, ** p<0.01, *** p<0.001; ns = not significant. (E) NMD *cis*-feature comparison of Cluster 5 subclusters vs. background (Bg): 3’UTR length, percentage of genes with an intron in the 3’UTR, and with a uORF; significance done by Wilcoxon rank-sum test for 3’UTR length, Fisher’s exact test for intron/uORF relative to background.

Cluster 5 (n = 279) displayed both genotype- and condition-specific trajectories. To separate these effects, cluster 5 was subdivided into three subclusters (subclusters 5/1–5/3) (Fig. 4B). Subcluster 5/1 and 5/3 genes, but not subcluster 5/2 genes, were overexpressed in the *upf3* mutant under all three conditions (NT, P/T and P/T+1d, respectively). Thus, NMD direct targets are mainly present in these subclusters, whereas subcluster 5/2 genes are predominantly indirect targets. Subcluster 5/1 was enriched for “RNA modification”, while subcluster 5/3 was weakly enriched for “defense response to symbiont” term (Fig. 4B,C). Subcluster 5/2 genes were expressed similarly at NT in the wild-type and *upf3*, remained unchanged in treated Col- 0, but were induced in *upf3* during heat (P/T) and further elevated upon recovery (P/T+1d). This subcluster was strongly enriched for immune-related terms, mirroring the genome-wide trends (Fig. 4C vs Fig S2D). These data suggest that the HS-inducible immune-related genes, which are suppressed by NMD, are mainly indirect NMD targets.

To distinguish direct transcriptional induction from NMD-dependent RNA destabilization, we analyzed promoter *cis*-elements alongside NMD *cis*-features (Fig 4D,E). The promoters of subcluster 5/2 were enriched for the W-box (OR = 1.89, adjusted p = 0.08) and heat-responsive HSE motifs (OR = 2.02, adjusted p < 0.05), whereas subcluster 5/1 was mildly enriched for the DRE/CRT motif (OR = 1.7)(Fig 4D). While subclusters 5/1 and 5/3 were enriched in genes having uORF NMD feature (28% and 21.3% vs 10% in background set; Fig. 4E, *right*), and subcluster 5/3 showed enrichment for 3′UTRs with introns (8,9% vs 4,3%, Fig 4E, *middle*), NMD features were underrepresented in subcluster 5/2 genes (Fig 4E, *left*). These results suggest that the immune genes present in subcluster 5/2 are upregulated mainly through transcriptional activation, possibly via both WRKY and HSF transcription factor activity, whereas transcripts in 5/1 and 5/3 may be stabilized in the absence of NMD activity, in agreement with their trajectories. Importantly, the activation of the immune subcluster 5/2 depended on heat: the *upf3*/Col-0 ratio was 1.1- at NT, 3.0- at P/T and 3.8-fold at P/T+1d, while the genes did not respond to heat in Col-0.

Collectively, clustering analysis revealed that heat induces a transcriptional immune response, which is suppressed by NMD in wild-type plants but occurs in the NMD-deficient mutant.

### Reduced *HSFA2* levels is associated with a dampened HSR in *upf3*

It was previously suggested that NMD regulates *HsfA2* isoforms’ stability (J. Liu et al., 2013; Sugio et al., 2009) and through this also thermomemory. For this reason, we examined selected hallmark transcripts of heat stress-memory (Fig. 5A). *HSFA2* dropped to ∼78% in *upf3* vs Col- 0 at P/T (BH-adjusted p = 0.06, n = 3), whereas the expression of memory factor *HsfA3* remained low at P/T in both genotypes. The memory HSP targets *sHSP18.2* and *sHSP22.0* were mildly induced, both were higher in Col-0 plants vs *upf3* (∼0.77- and 0.65-fold decrease, respectively). The control non-memory *HSP101* was induced at P/T, non-significantly lower in *upf3* (∼88% of Col-0). During recovery the level of *HsfA3* and memory chaperons sHSP18.2 and sHSP22 was strongly elevated, with no significant difference between genotypes (Fig 5A). To examine the HSR genome-widely, we defined heat-responsive genes as those induced in Col-0 relative to NT (HSR genes, n = 2416, BH-adjusted p < 0.05, log_2_FC > 1), and analyzed the difference in HSR expression between Col-0 and *upf3*, using the non-HSR set (n = 17,711) as control. Importantly, the mean expression of the HSR set was significantly higher in *upf3* relative to Col-0 at NT (∼1.27-fold), but significantly lower at P/T (0.86-fold) (Fig 5B). The difference disappeared at P/T+1d. These results suggest that NMD-deficiency (i) already creates a weak heat-stress-primed state under NT conditions, and (ii) dampens the response during the acute phase of heat stress, an effect that is mostly lost during recovery. The findings that both HSFA2 induction and the HSR activation are weaker in the NMD-deficient mutant at P/T suggest that NMD is required for optimal HSFA2 mRNA level and HS responses.

**Figure 5.**
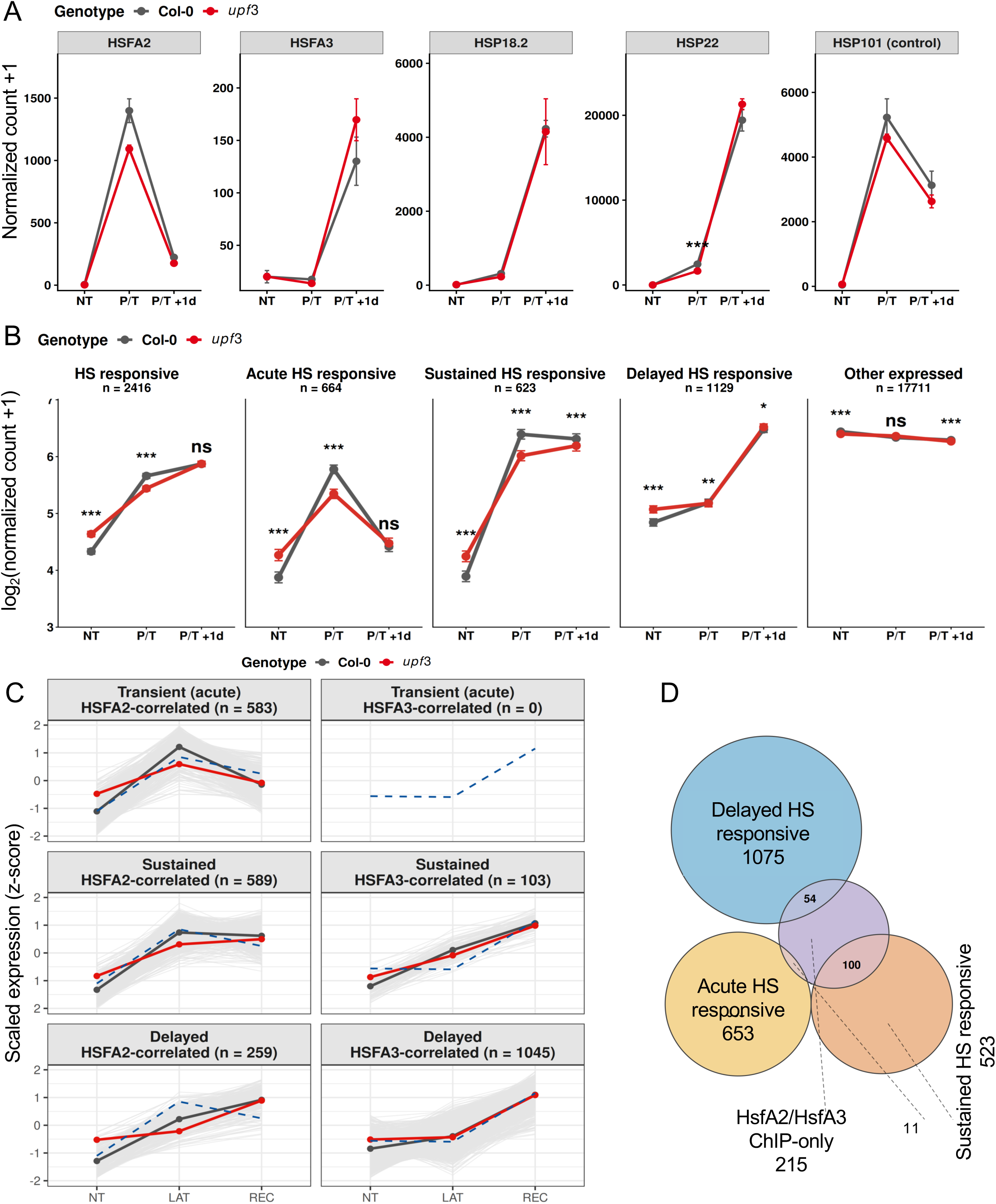
NMD loss in *upf3* downturns the HsfA2-, but not HsfA3-driven, HS-memory program. (A) Normalized count numbers of the heat-stress memory factors HsfA2 and HsfA3, and their targets sHSP18.2 and HSP22 (group means, ± SE, n=3; hypergeometric test, BH-adjp ***p<0.001). (B) Estimated mean from a genome-wide mixed-effects model for the all HS responsive, transient, sustained and delayed HSR genes vs. the non-HSR control, hypergeometric test, BH-adjp ***p<0.001, non-significant, ns; (C) Mean scaled expression trajectories of the three HSR groups with HsfA2 and HsfA3 trajectories (blue dashed lines) at NT, P/T and REC; WT transcripts (grey), *upf3* (red), thin lines represent single gene trajectories; (D) Overlap between the HSR gene groups and the combined HsfA2/HsfA3 ChIP-seq target gene set (Kappel et al., 2023), gene numbers are shown.

Although both HSFA2 and HSFA3 are essential for thermomemory, our findings that HSFA2 is upregulated at P/T and decreases at P/T+1d, while HSFA3 is induced at P/T+1d indicate that they could regulate the HSR at different stages. Based on these, we grouped the heat-responsive genes into transient (P/T-specific, n = 664), sustained (increased level during both P/T and P/T+1d, n = 623) and delayed (P/T+1d-specific, n = 1,129) sets (Fig. 5B). We found the transient genes are enriched in cluster 6, the sustained genes in cluster 2 and mildly cluster 6, while the delayed genes in cluster 4 (Fig. S4). Expression of most transient and sustained genes correlated with *HsfA2* (n = 583, 93.6%, and n = 589, 94.5%, respectively); while the delayed genes correlated mainly with *HsfA3* (n = 1045) (Fig 5C, S5A). Relevantly, the largest genotype differences were seen in the HSFA2-correlated transient and sustained sets at P/T (∼0.74- and 0.77-fold decrease in *upf3*), and the smallest in the HsfA3-correlated delayed set at both P/T and P/T+1d (∼0.99-fold decrease in *upf3*) (Fig 5C), indicating that specifically HsfA2 pathway is affected by NMD loss.

The direct target loci of HsfA2/HsfA3 were defined by ChIP-seq (Kappel et al., 2023): the two factors bind to nearly identical loci, in total 380 genes. We analyzed the overlap of our three HSR groups with these ChIP loci: 11 of 664 transient genes (1.7%), 100 of 623 sustained genes (16%) and 54 of 1,129 delayed genes (4.8%) overlapped with HsfA2/A3-ChIP loci (Fig. 5D). Of the 215 ‘ChIP-only’ genes left, 94 showed no significant change, 65 were weakly induced, 30 were not or only lowly expressed and 26 were repressed by heat in our dataset (Fig. S4B); overall, ChIP-only genes had a lower expression level in *upf3* (Fig. S4C) and were enriched for GO terms “nucleotide catabolic processes” and weakly for “response to heat” (Fig S4D). Notably, the transient and sustained HSR genes were found to be significantly lower expressed at P/T in *upf3*, but no difference was found in the delayed genes between genotypes (Fig 5C). The strongest induction occurred in the ChIP-bound (much less in the unbound) sustained genes, at both P/T and P/T+1d, with lower median in *upf3* (Fig. S4E).

To test whether the lower transcript levels in *upf3* plants reflect transcription or decay, we analyzed promoter and NMD *cis*-features (Fig S5B,C). For the transient set we found promoter enrichment of HSE (OR = 1.26, adjp < 0.05), in case of the sustained set for HSE (OR = 1.25, adjp < 0.05) and ABRE (OR = 1.79, adjp < 0.001), and for the delayed set of ABRE (OR = 1.31, adjp < 0.01) and DRE (OR = 1.21, adjp < 0.05) *cis* elements (Fig S5B). In addition, no significant enrichment of genes with NMD features could be found in the three HSR sets (Fig. S5C). These findings are consistent with HSR genes being transcriptionally regulated through HSE (transient set), HSE and ABRE (sustained set), and ABRE and DRE (delayed set) *cis* elements within their promoters and being rarely direct NMD targets.

Based on these results, we propose that the transient and sustained sets of HSR genes are regulated primarily by HsfA2, while the delayed HSR set is controlled by HsfA3, and that, the decreased level of *HsfA2* mRNA in *upf3* leads to the drop of transient and sustained genes at P/T.

### NMD deficiency leads to HsfA2- and HSFA3-independent early heat tolerance

The transcriptome analysis showed a reduced level of HSFA2 mRNA in *upf3* and a dampened induction of the HsfA2-correlated, but not the HsfA3-correlated branch (Fig. 5). This may explain the ineffective late recovery of *upf3*, but not the early heat tolerance of the NMD mutants (Fig. 1, S1). We therefore hypothesized that NMD can also act independently of the HsfA2/HsfA3 pathways and tested this using the double mutants *upf3-1;hsfa2-1* (*upf3;hsfa2*) and *upf3-1;hsfa3-1* (*upf3;hsfa3*). Under NT conditions, *upf3* rosettes were slightly smaller than those of Col-0 (Arciga-Reyes et al., 2006), whereas *hsfa2* and *hsfa3* rosettes appeared slightly larger. *hsfa2* plants had started to bolt earlier than *Col-0, upf3* and *hsfa3*. The *upf3;hsfa3* rosettes resembled *upf3* in size, whereas *upf3;hsfa2* rosettes were larger and were also bolting (Fig. S6).

First, we analyzed NMD activity in the *upf3* and *hsfa2* single and the *upf3,hsfa2* double mutants using the *SMG7* (Z. Kerenyi et al., 2008) and *At5g45430* (Degtiar et al., 2015) NMD reporters. Both NMD-sensitive transcripts were elevated in the *upf3* and *upf3,hsfa2* genotypes under all conditions and were unchanged in the *hsfa2* mutant, indicating that NMD is not affected by the *hsfa2* mutation (Fig. S7).

Further, we compared the heat sensitivity to P/T treatments in a time series treatment set (50, 60 and 80 min; Fig. 6, S8). The *hsfa2* and *hsfa3* single mutants were markedly heat-sensitive (Fig. 6A,B; S8A,B), confirming the roles of HsfA2 and HsfA3 in thermomemory. During early recovery (P/T+4d), *upf3* was less affected than the Col-0. The double mutants were less sensitive than the respective *hsfa* single mutants but remained more sensitive than the *upf3* (Fig. 6B, S8B). This suggests that the loss of UPF3 improves early heat tolerance likely independently of HsfA2 and HsfA3. At late recovery (P/T+14d), the *upf3;hsfa2* and *upf3;hsfa3* plants were not significantly different from *hsfa2* and *hsfa3* (Fig. 6C,D, S8C,D), thus, the loss of NMD could not compensate for the high mortality caused by the *hsfa2* and *hsfa3* mutations.

**Figure 6:**
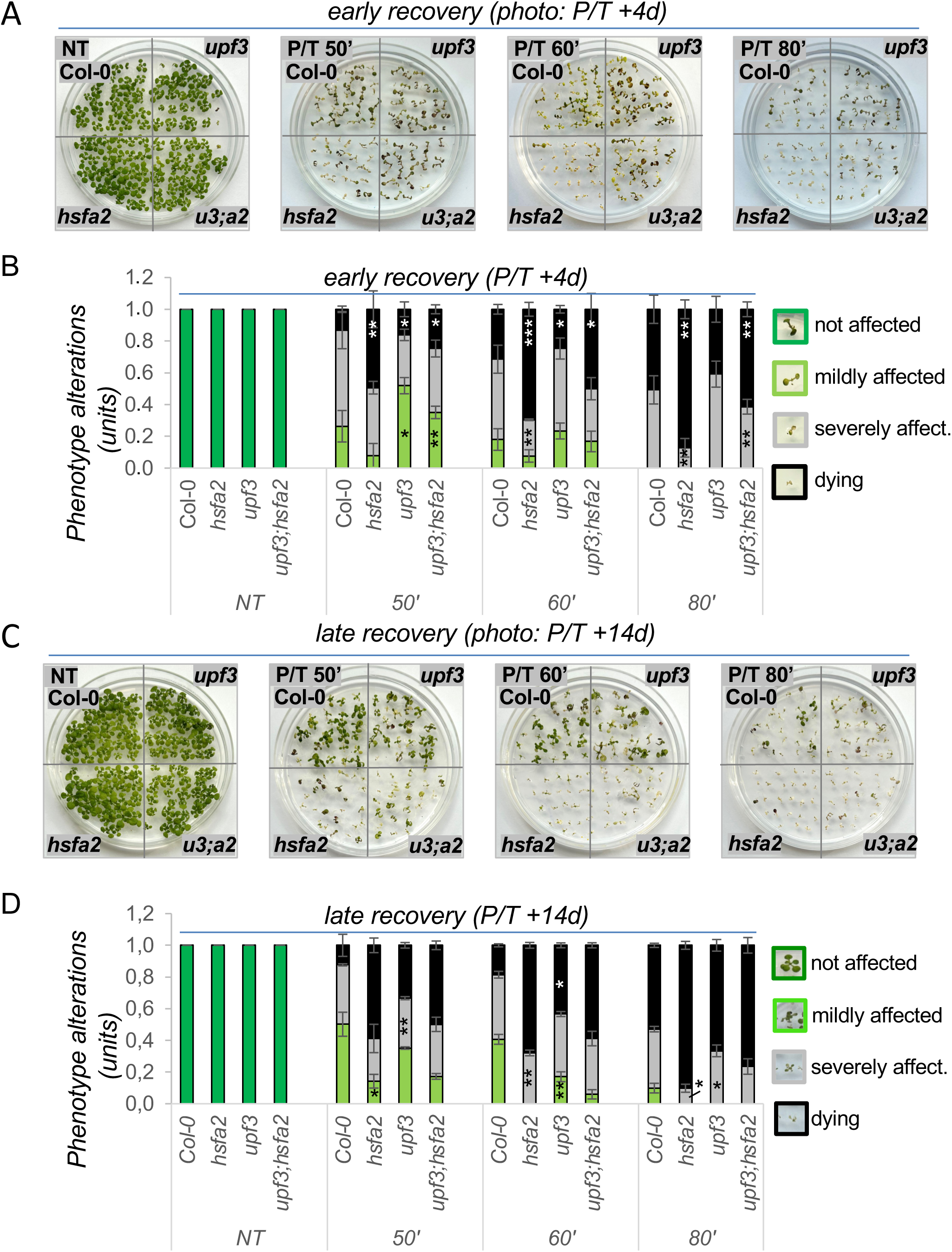
NMD’s impact on heat-stress recovery is independent of the HsfA2 pathway. (A) Heat-stress phenotypes of wild-type (Col-0), *upf3*, *hsfa2*, and *upf3;hsfa2* (*u3;a2*) mutants on day 4 post-treatment (P/T+4d, early recovery); (B) Quantification of early-recovery phenotypes shown in *panel A*; (C) phenotypes of the same genotypes on day 14 post-treatment (P/T+14d, late recovery). (D) quantification of late-recovery phenotypes shown in *panel B*; data are mean ± SD of three independent experiments, and asterisks indicate significant differences of single mutants vs. Col-0 or of the double mutant vs. *hsfa2* (*p < 0.05; **p < 0.01; ***p < 0.001, unpaired two-sided t-test).

Together, these results suggest that UPF3 loss confers a HSFA2- and HSFA3-independent, transient heat tolerance during early recovery, whereas HsfA2 and HsfA3 remain the main drivers of long-term memory.

### NMD deficiency confers early heat tolerance independently of the HsfA2 regulon

Several HSFs (HsfA4c, HsfA7b, HsfB1 and HsfB2a) were previously proposed to be NMD targets (J. Liu et al., 2013). To test whether the altered heat tolerance of the NMD mutants is based on the altered priming response of these NMD targets, we measured the amount of these transcripts by RT-qPCR at NT, P and P+1d (Fig S9). Surprisingly, none of them was a genuine NMD target, they were not upregulated in the *upf3* mutant at NT. Moreover, their expression changed similarly in the Col-0 and *upf3* plants during the treatments, indicating that these HSFs, which were previously proposed to be NMD-regulated, were not responsible for the early heat tolerance of NMD mutant (Fig S9).

Previously we observed that HSR transcripts are upregulated in *upf3* already at NT condition (Fig 5B,C). These could contribute to the early heat tolerance of *upf3* plants. To evaluate the extent to which these mRNAs confer an overall stress-primed status to NMD deficient plants, we calculated a priming index (PI, 1680 out of the total 2358 HS-responsive transcripts were significantly induced in *upf3*): the *upf3*/Col-0 log_2_FC values at NT (genotype difference) was divided by the maximum WT induction at P/T (condition difference). In this ratio the PI=1 means full heat stress level of the given transcript. Across all 1,680 HSR genes the median PI was 22% in *upf3* plants (Fig S10). When the PI was calculated on ChIP-bound vs -unbound sets to assess the HsfA2/A3 impact: the PI for the ChIP-bound group was almost zero. This shows that the effect was entirely confined to genes not bound by HsfA2/A3 (35%, 30% and 18%; ChIP-bound groups −21%, −2% and −7%). These results suggest that the mild enhanced HSR expression in the NMD mutant at NT is not caused by HsfA2/A3 binding and induction (Fig S10A). Further, removing 49 known NMD targets (Fig S10A, *middle panel*), or 277 genes with known or predicted NMD structural features (Fig S10. *bottom panel*) changed the PI median only modestly (all HSR classes 22%, 22%, 20%; transient 34%, 33%, 33%; sustained 22%, 22%, 19%; delayed 16%, 15%, 14%), suggesting that the primed state is mostly an indirect effect of NMD loss.

To try to define the nature of this priming, we next analyzed the GO enrichment of the HSR genes already elevated in *upf3* at NT (n = 1680). No typical heat-related terms were enriched. Instead, “RNA modification” was the only term enriched, mirroring the genome-wide GO analysis (Fig. S10B). This suggests that the stress priming status of *upf3* under ambient conditions (PI ∼22%) is achieved through non-canonical pathways, namely RNA modification. This may transiently improve the heat tolerance of NMD mutants during early recovery, although this protection does not seem to be lasting on longer time periods (Fig. 6D).

### NMD-controlled *HsfA2-II* and *HsfA2-III* isoforms may indirectly lower *HsfA2-I* abundance

NMD has previously been proposed to influence heat stress memory through the *HSFA2* splice isoforms *HsfA2-II* and *HsfA2-III*, of which *HsfA2-III* acts as a positive regulator of HSFA2 transcription (J. Liu et al., 2013; Sugio et al., 2009). Our RNA-seq data showed that the overall *HSFA2* transcript level is lower in *upf3* at P/T (Fig. 5A). To see how the signal of the HSFA2 locus is distributed between isoforms, we measured the functional, non-NMD-target isoform *HsfA2-I* and the NMD-target isoforms *HsfA2-II* and *HsfA2-III* separately by RT-qPCR after priming (P) in Col-0, *upf3*, *hsfa2* and the double mutant *upf3;hsfa2* (Fig. 7A–C). All three isoforms were induced at P in Col-0 and *upf3* and returned to basal levels at P+1d. None of them were detected in *hsfa2* or *upf3;hsfa2*, confirming their mutant status. Compared with Col- 0, *upf3* had ∼35% less *HsfA2-I* but about 6-fold more *HsfA2-II*. *HsfA2-III* showed the same trend (∼2-2.5-fold increase). The accumulation of *HsfA2-II* and *HsfA2-III* in *upf3* is consistent with their degradation by NMD (J. Liu et al., 2013; Sugio et al., 2009).

**Figure 7:**
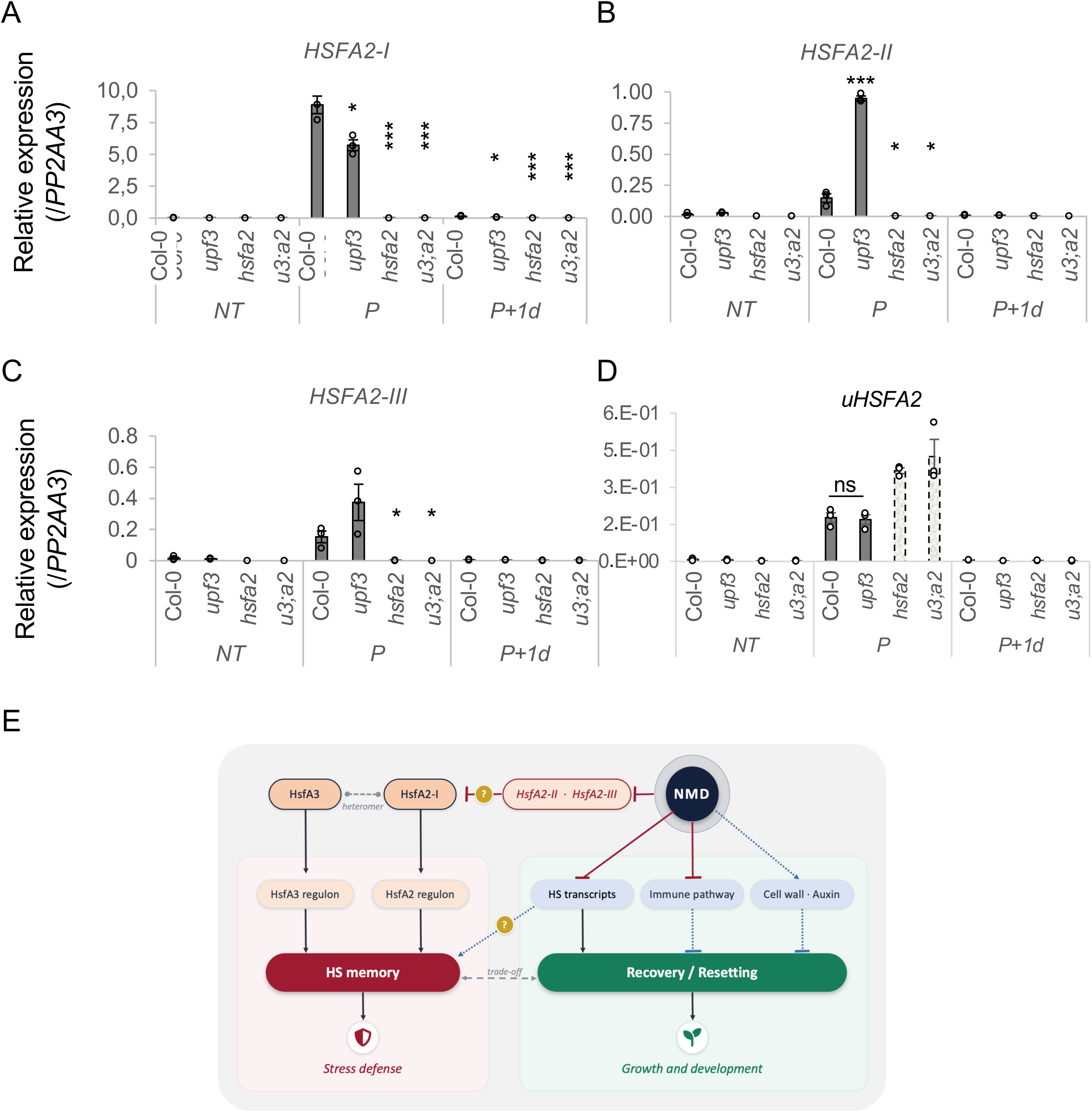
NMD modulates HSR through HsfA2-independent and -dependent routes. RT-qPCR of (A) *HsfA2-I*, (B) *HsfA2-II*, (C) *HsfA2-III* and (D) unspliced *HsfA2* (*uHsfA2*) transcripts in Col-0, *upf3*, *hsfa2* and *upf3;hsfa2* (*u3;a2*) plants that were non-treated (NT), sampled after priming (P), or one day later (P+1d); expression is relative to *PP2AA3*. Data are mean ± SD of at least three biological replicates; P-values are from two-tailed Student’s t-tests for differences relative to Col-0 within each condition (*P < 0.05, **P < 0.01, ***P < 0.001; ns, not significant). In (D), dashed bars (*hsfa2* and *u3;a2*) indicate signal most likely arising from a chimeric, non-functional *hsfa2* T-DNA transcript, since the primers lie upstream of the T-DNA insertion. (E) Working model: black arrows debote activation, red bars inhibition, dotted blue arrows/bars indirect effects, and “?” untested links; for details see main text.

*HsfA2-I* is not an NMD target, so its decrease in *upf3* must be indirect. One possible explanation is feedback on the transcription of the locus. To test this, we measured the unspliced *HSFA2* transcript (*uHSFA2*) as a proxy for transcriptional activity. The *uHSFA2* levels did not differ between Col-0 and *upf3* (Fig. 7D; note, the signal in *hsfa2* and *upf3;hsfa2* most likely arises from a chimeric T-DNA transcript), arguing against reduced transcription. We therefore propose that, with unchanged transcription, a shift in splicing from *HsfA2-I* toward the NMD- controlled isoforms *HsfA2-II/III* may indirectly reduce *HsfA2-I* abundance in *upf3* (Fig. 7E).

## Discussion

The heat shock response (HSR) is a multilayered process in which several regulatory mechanisms act in parallel, including changes in RNA and protein homeostasis (Chung et al., 2025; Li, Tang, Srivastava, Bassham, & Howell, 2020; Rahaman et al., 2025; Staacke, Mueller- Roeber, & Balazadeh, 2025). RNA decay is one of these layers. The exoribonuclease XRN4 degrades transcripts that are no longer needed (Merret et al., 2013; Merret et al., 2015; Nguyen et al., 2015) whereas NMD is a surveillance pathway that safeguards transcript fidelity by eliminating erroneous transcripts (Szaker et al., 2025). Moreover, NMD also directly targets normal mRNAs possessing NMD inducing features and indirectly controls the genes that are regulated by these direct targets. Timely RNA decay facilitates efficient transcriptome reprogramming and releases cellular resources for growth and recovery.

Building on our previous work (Szaker et al., 2025), we examined how NMD contributes to the heat stress recovery in *A. thaliana*, and tested whether it regulates the HsfA2 pathway, a link previously proposed between NMD and thermomemory (J. Liu et al., 2013; Sugio et al., 2009). By comparing the thermotolerance of wild-type plants with the *nmd* single and *nmd;hsfa2* and *nmd;hsfa3* double mutants in LAT assays, we showed that NMD plays a dual role in thermotolerance regulation; it inhibits short-term heat tolerance in a HSFA2/3- independent manner, whereas it stimulates long-term HS tolerance. To reveal the molecular basis of this dual role, transcriptomes of wild-type and NMD mutants were compared at the NT, P/T and P/T+1d timepoints. We found that NMD was active throughout the LAT assay (Fig 3). Transcriptome analysis of NMD-deficient mutants revealed four main features. First, more genes were upregulated than downregulated in the NMD mutant, and many of the upregulated transcripts carry direct NMD features (Fig 3, S2). Second, NMD deficiency creates a stress-primed state in NT plants through the upregulation of HsfA2-independent transcripts involved in RNA modification and HSR (Fig 5B,C, S10). We propose that this stress-primed state explains the early, transient thermotolerance; although the mechanism remains to be determined. Third, HSFA2 activity is reduced in the *upf3* plant, *HSFA2-I* mRNA accumulates to a lower level and the balance between functional and non-functional HsfA2 isoforms is shifted at P/T (Fig 7). We propose that this low *HSFA2-I* level leads to the diminished induction of HSFA2-dependent memory genes; they accumulate to about 80% of wild-type levels (Fig. 5). Fourth, NMD deficiency shifted the transcriptome toward immune signaling at the expense of auxin-related growth and cell-wall biogenesis during HS and recovery (Fig 4, S2). Because maintaining a stress state is metabolically costly (Cadena-Ramos, Rodriguez-Pina, & De-la- Pena, 2026; Staacke et al., 2025; Zhou et al., 2013), this altered growth–defense balance and the weakened memory regulon may explain the physiological decline (reduced chlorophyll, lower photosynthetic efficiency, higher ion leakage, Fig 2) and late heat sensitivity of NMD mutants (Fig 1, 6, S1, S8).

The elevated expression of immune genes in *upf3*, which appears mainly when plants are exposed to heat, suggests that NMD restrains defense responses during heat stress, which may allow plants to shift resources from defense to regrowth. This trade-off could matter in nature when heat waves are followed by pathogen infections. Infection, in contrast, inhibits NMD, in part by reducing the levels of UPF proteins (Gloggnitzer et al., 2014), whereas heat does not reduce NMD efficiency (Fig. 5). Whether this crosstalk between heat and biotic stress is biologically relevant remains to be tested, e.g. by generating NMD and immune double mutants, and then infecting them upon/after heat treatment.

Our data indicate that the NMD-dependent regulation of HsfA2 expression and isoform imbalance affects the HsfA2-regulon, but we detected no effect on the HsfA3-regulon. Since HSFA2/HSFA3 heteromers promote HS memory (Friedrich et al., 2021), NMD-mediated control of HsfA2 plays a modulatory role in thermomemory. HsfA2 and HsfA3 share their targets (Kappel et al., 2023) but differ in kinetics (HsfA2 is strongly induced at P/T; HsfA3 is mildly elevated at P/T and strongly at P/T+1d). The NMD-dependent modulation of HsfA2 may finetune the temporal composition of HSF complexes, and optimize the HS responses. The S-HsfA2 peptide translated from *HsfA2-III* binds HREs, thereby dampening the induction of HS-responsive genes, and attenuating full-length HsfA2 activity (Chen et al., 2025). The stabilized *HsfA2-III* in *upf3* may therefore contribute to the downregulation of the HS regulon. However, this cannot explain the isoform changes in *upf3* (*HsfA2-I* ∼35% down, *HsfA2-II* ∼6- fold up, *HsfA2-III* ∼2.2-fold up, *uHsfA2* unchanged). The unchanged expression of the *uHsfA2* transcripts excludes altered transcription, so the effect must be post-transcriptional. NMD acts downstream of splicing and should destabilize *HsfA2-II/III* without affecting *HsfA2-I*. Non- mutually exclusive feedback regulatory routes could include competition for splicing factors, altered nuclear export and positive regulation of unproductive splicing. Regulation of a functional isoform by an NMD-target isoform may be a widespread phenomenon and deserves further study.

In summary, NMD functions as a key regulator of thermal acclimation. By destabilizing stress- related transcripts under ambient conditions, constraining immune gene overexpression during both ambient and heat stress, and fine-tuning the functional isoforms of HsfA2 (and consequently the HsfA2 thermomemory pathway), NMD operates as an RNA quality control pathway to orchestrate the transition from a stress response to physiological resetting/regrowth. Together with recent evidence, these findings support a model in which NMD is a fundamental mechanism for maintaining transcriptome integrity in fluctuating environments.

## Author contribution

SH Abbas: genotyping, crosses, heat stress treatments, phenotyping, molecular measurements, data analysis, writing; HM Szaker performed bioinformatic and data analysis; E Darkó: physiological assays and data analysis, T Erős: genotyping, HS-phenotyping; D Silhavy: data analysis, funding, writing, T Csorba: conceptualization, funding, writing. All authors contributed to the final version.

## Data availability

Data and materials are available upon request.

## References

Akerfelt, M., Morimoto, R. I., & Sistonen, L. (2010). Heat shock factors: integrators of cell stress, development and lifespan. Nat Rev Mol Cell Biol, 11(8), 545–555. doi:10.1038/nrm2938

Asghar, M.A., Balogh, E., Szalai, G., Galiba, G., Kocsy, G. (2022). Differences in the light-dependent changes of the glutathione metabolism during cold acclimation in wheat varieties with different freezing tolerance. Journal of Agronomy and Crop Science 208(1). 65–75. doi: 10.1111/jac.12566

Arciga-Reyes, L., Wootton, L., Kieffer, M., & Davies, B. (2006). UPF1 is required for nonsense-mediated mRNA decay (NMD) and RNAi in Arabidopsis. Plant J, 47(3), 480–489. doi:10.1111/j.1365-313X.2006.02802.x

Cadena-Ramos, A. I., Rodriguez-Pina, A. L., & De-la-Pena, C. (2026). ROS and autophagy in plant stress responses: adaptive partners in survival. Plant Physiol Biochem, 231, 110991. doi:10.1016/j.plaphy.2025.110991

Charng, Y. Y., Liu, H. C., Liu, N. Y., Chi, W. T., Wang, C. N., Chang, S. H., & Wang, T. T. (2007). A heat-inducible transcription factor, HsfA2, is required for extension of acquired thermotolerance in Arabidopsis. Plant Physiol, 143(1), 251–262. doi:10.1104/pp.106.091322

Charng, Y. Y., Mitra, S., & Yu, S. J. (2023). Maintenance of abiotic stress memory in plants: Lessons learned from heat acclimation. Plant Cell, 35(1), 187–200. doi:10.1093/plcell/koac313

Chen, W., Zhao, J., Tao, Z., Zhang, S., Bei, X., Lu, W., & Qi, X. (2025). Short heat shock factor A2 regulates heat resistance and growth balance in Arabidopsis. Elife, 13. doi:10.7554/eLife.99937

Chung, K. K., Law, K. C., Zhao, Z., Ma, J., Zhan, X. T., Chiang, C. H., … Zhuang, X. (2025). Biomolecular condensation of ERC1 recruits ATG8 and NBR1 to drive autophagosome formation for plant heat tolerance. Proc Natl Acad Sci U S A, 122(46), e2425689122. doi:10.1073/pnas.2425689122

Danecek, P., Bonfield, J. K., Liddle, J., Marshall, J., Ohan, V., Pollard, M. O., … Li, H. (2021). Twelve years of SAMtools and BCFtools. Gigascience, 10(2). doi:10.1093/gigascience/giab008

Darkó, É., Fodor, J., Dulai, S., Ambrus, H., Szenzenstein, A., Király, Z., Barnabás, B. (2011). Improved Cold and Drought Tolerance of Doubled Haploid Maize Plants Selected for Resistance to Prooxidant tert-Butyl Hydroperoxide. Journal of agronomy and crop science 197(6), 454–465. doi: 10.1111/j.1439-037X.2011.00479.x

Degtiar, E., Fridman, A., Gottlieb, D., Vexler, K., Berezin, I., Farhi, R., … Shaul, O. (2015). The feedback control of UPF3 is crucial for RNA surveillance in plants. Nucleic Acids Res, 43(8), 4219–4235. doi:10.1093/nar/gkv237

Friedrich, T., Oberkofler, V., Trindade, I., Altmann, S., Brzezinka, K., Lamke, J., … Baurle, I. (2021). Heteromeric HSFA2/HSFA3 complexes drive transcriptional memory after heat stress in Arabidopsis. Nat Commun, 12(1), 3426. doi:10.1038/s41467-021-23786-6

Gloggnitzer, J., Akimcheva, S., Srinivasan, A., Kusenda, B., Riehs, N., Stampfl, H., … Riha, K. (2014). Nonsense-mediated mRNA decay modulates immune receptor levels to regulate plant antibacterial defense. Cell Host Microbe, 16(3), 376–390. doi:10.1016/j.chom.2014.08.010

Kappel, C., Friedrich, T., Oberkofler, V., Jiang, L., Crawford, T., Lenhard, M., & Baurle, I. (2023). Genomic and epigenomic determinants of heat stress-induced transcriptional memory in Arabidopsis. Genome Biol, 24(1), 129. doi:10.1186/s13059-023-02970-5

Karanyi, Z., Mosolygo, L. A., Fero, O., Horvath, A., Boros-Olah, B., Nagy, E., … Szekvolgyi, L. (2022). NODULIN HOMEOBOX is required for heterochromatin homeostasis in Arabidopsis. Nat Commun, 13(1), 5058. doi:10.1038/s41467-022-32709-y

Kerenyi, F., Wawer, I., Sikorski, P. J., Kufel, J., & Silhavy, D. (2013). Phosphorylation of the N- and C-terminal UPF1 domains plays a critical role in plant nonsense-mediated mRNA decay. Plant J, 76(5), 836–848. doi:10.1111/tpj.12346

Kerenyi, Z., Merai, Z., Hiripi, L., Benkovics, A., Gyula, P., Lacomme, C., … Silhavy, D. (2008). Inter-kingdom conservation of mechanism of nonsense-mediated mRNA decay. EMBO J, 27(11), 1585–1595. doi:10.1038/emboj.2008.88

Kertesz, S., Kerenyi, Z., Merai, Z., Bartos, I., Palfy, T., Barta, E., & Silhavy, D. (2006). Both introns and long 3’-UTRs operate as cis-acting elements to trigger nonsense-mediated decay in plants. Nucleic Acids Res, 34(21), 6147–6157. doi:10.1093/nar/gkl737

Kurihara, Y., Matsui, A., Hanada, K., Kawashima, M., Ishida, J., Morosawa, T., … Seki, M. (2009). Genome-wide suppression of aberrant mRNA-like noncoding RNAs by NMD in Arabidopsis. Proc Natl Acad Sci U S A, 106(7), 2453–2458. doi:10.1073/pnas.0808902106

Li, Z., Srivastava, R., Howell, S. H., Chudalayandi, S., & Severin, A. (2026). The Role of Nonsense-Mediated Decay in Alleviating Nonproductive RNAs Produced by Heat Stress-Induced Alternative Splicing in Arabidopsis. Plant Cell Environ, 49(3), 1561–1565. doi:10.1111/pce.70321

Li, Z., Tang, J., Srivastava, R., Bassham, D. C., & Howell, S. H. (2020). The Transcription Factor bZIP60 Links the Unfolded Protein Response to the Heat Stress Response in Maize. Plant Cell, 32(11), 3559–3575. doi:10.1105/tpc.20.00260

Liao, Y., Smyth, G. K., & Shi, W. (2014). featureCounts: an efficient general purpose program for assigning sequence reads to genomic features. Bioinformatics, 30(7), 923–930. doi:10.1093/bioinformatics/btt656

Lichtenthaler, H. K. (1987). Chlorophylls and carotenoids: Pigments of photosynthetic biomembranes. Methods in Enzymology 148, 350–382. doi: 10.1016/0076-6879(87)48036-1

Liu, H. C., & Charng, Y. Y. (2013). Common and distinct functions of Arabidopsis class A1 and A2 heat shock factors in diverse abiotic stress responses and development. Plant Physiol, 163(1), 276–290. doi:10.1104/pp.113.221168

Liu, J., Sun, N., Liu, M., Liu, J., Du, B., Wang, X., & Qi, X. (2013). An autoregulatory loop controlling Arabidopsis HsfA2 expression: role of heat shock-induced alternative splicing. Plant Physiol, 162(1), 512–521. doi:10.1104/pp.112.205864

Lloyd, J. P., & Davies, B. (2013). SMG1 is an ancient nonsense-mediated mRNA decay effector. Plant J, 76(5), 800–810. doi:10.1111/tpj.12329

Lloyd, J. P. B. (2018). The evolution and diversity of the nonsense-mediated mRNA decay pathway. F1000Res, 7, 1299. doi:10.12688/f1000research.15872.2

Love, M. I., Huber, W., & Anders, S. (2014). Moderated estimation of fold change and dispersion for RNA-seq data with DESeq2. Genome Biol, 15(12), 550. doi:10.1186/s13059-014-0550-8

Luha, R., Rana, V., Vainstein, A., & Kumar, V. (2024). Nonsense-mediated mRNA decay pathway in plants under stress: general gene regulatory mechanism and advances. Planta, 259(3), 51. doi:10.1007/s00425-023-04317-7

Martin, M. (2011). Cutadapt removes adapter sequences from high-throughput sequencing reads. EMBnet J. 17(1), doi: 10.14806/ej.17.1.200

Merret, R., Descombin, J., Juan, Y. T., Favory, J. J., Carpentier, M. C., Chaparro, C., … Bousquet-Antonelli, C. (2013). XRN4 and LARP1 are required for a heat-triggered mRNA decay pathway involved in plant acclimation and survival during thermal stress. Cell Rep, 5(5), 1279–1293. doi:10.1016/j.celrep.2013.11.019

Merret, R., Nagarajan, V. K., Carpentier, M. C., Park, S., Favory, J. J., Descombin, J., … Bousquet-Antonelli, C. (2015). Heat-induced ribosome pausing triggers mRNA co-translational decay in Arabidopsis thaliana. Nucleic Acids Res, 43(8), 4121–4132. doi:10.1093/nar/gkv234

Nasim, Z., Karim, N., Blilou, I., & Ahn, J. H. (2025). NMD-mediated posttranscriptional regulation fine-tunes the NLR-WRKY regulatory module to modulate bacterial defense response. Plant Sci, 356, 112528. doi:10.1016/j.plantsci.2025.112528

Nguyen, A. H., Matsui, A., Tanaka, M., Mizunashi, K., Nakaminami, K., Hayashi, M., … Seki, M. (2015). Loss of Arabidopsis 5’-3’ Exoribonuclease AtXRN4 Function Enhances Heat Stress Tolerance of Plants Subjected to Severe Heat Stress. Plant Cell Physiol, 56(9), 1762–1772. doi:10.1093/pcp/pcv096

Obermeyer, S., Stockl, R., Schnekenburger, T., Kapoor, H., Stempfl, T., Schwartz, U., & Grasser, K. D. (2023). TFIIS Is Crucial During Early Transcript Elongation for Transcriptional Reprogramming in Response to Heat Stress. J Mol Biol, 435(2), 167917. doi:10.1016/j.jmb.2022.167917

Ohama, N., Sato, H., Shinozaki, K., & Yamaguchi-Shinozaki, K. (2017). Transcriptional Regulatory Network of Plant Heat Stress Response. Trends Plant Sci, 22(1), 53–65. doi:10.1016/j.tplants.2016.08.015

Ohtani, M., & Wachter, A. (2019). NMD-Based Gene Regulation-A Strategy for Fitness Enhancement in Plants? Plant Cell Physiol, 60(9), 1953–1960. doi:10.1093/pcp/pcz090

Rahaman, S., Schiffelholz, N., Mittal, N., Frohlich, K. E., Zavolan, M., & Becskei, A. (2025). Heat shock induces silent ribosomes and reorganizes mRNA turnover. Cell Rep, 44(10), 116447. doi:10.1016/j.celrep.2025.116447

Riehs-Kearnan, N., Gloggnitzer, J., Dekrout, B., Jonak, C., & Riha, K. (2012). Aberrant growth and lethality of Arabidopsis deficient in nonsense-mediated RNA decay factors is caused by autoimmune-like response. Nucleic Acids Res, 40(12), 5615–5624. doi:10.1093/nar/gks195

Staacke, T., Mueller-Roeber, B., & Balazadeh, S. (2025). Stress resilience in plants: the complex interplay between heat stress memory and resetting. New Phytol, 245(6), 2402–2421. doi:10.1111/nph.20377

Sugio, A., Dreos, R., Aparicio, F., & Maule, A. J. (2009). The cytosolic protein response as a subcomponent of the wider heat shock response in Arabidopsis. Plant Cell, 21(2), 642–654. doi:10.1105/tpc.108.062596

Szadeczky-Kardoss, I., Szaker, H. M., Verma, R., Darko, E., Pettko-Szandtner, A., Silhavy, D., & Csorba, T. (2022). Elongation factor TFIIS is essential for heat stress adaptation in plants. Nucleic Acids Res, 50(4), 1927–1950. doi:10.1093/nar/gkac020

Szaker, H. M., Darko, E., Medzihradszky, A., Janda, T., Liu, H. C., Charng, Y. Y., & Csorba, T. (2019). miR824/AGAMOUS-LIKE16 Module Integrates Recurring Environmental Heat Stress Changes to Fine-Tune Poststress Development. Front Plant Sci, 10, 1454. doi:10.3389/fpls.2019.01454

Szaker, H. M., Verma, R., Szadeczky-Kardoss, I., Gal, N., Abbas, S. H., Darko, E., … Csorba, T. (2025). Transcription fidelity and control of alternative splicing contribute to heat stress survival in Arabidopsis. Plant Cell, 37(11). doi:10.1093/plcell/koaf256

Weng, M., Yang, Y., Feng, H., Pan, Z., Shen, W. H., Zhu, Y., & Dong, A. (2014). Histone chaperone ASF1 is involved in gene transcription activation in response to heat stress in Arabidopsis thaliana. Plant Cell Environ, 37(9), 2128–2138. doi:10.1111/pce.12299

Wu, H. L., Ai, Q., Teixeira, R. T., Nguyen, P. H. T., Song, G., Montes, C., … Hsu, P. Y. (2024). Improved super-resolution ribosome profiling reveals prevalent translation of upstream ORFs and small ORFs in Arabidopsis. Plant Cell, 36(3), 510–539. doi:10.1093/plcell/koad290

Wu, T., Hu, E., Xu, S., Chen, M., Guo, P., Dai, Z., … Yu, G. (2021). clusterProfiler 4.0: A universal enrichment tool for interpreting omics data. Innovation (Camb), 2(3), 100141. doi:10.1016/j.xinn.2021.100141

Yoshida, T., Ohama, N., Nakajima, J., Kidokoro, S., Mizoi, J., Nakashima, K., … Yamaguchi-Shinozaki, K. (2011). Arabidopsis HsfA1 transcription factors function as the main positive regulators in heat shock-responsive gene expression. Mol Genet Genomics, 286(5-6), 321–332. doi:10.1007/s00438-011-0647-7

Zhou, J., Wang, J., Cheng, Y., Chi, Y. J., Fan, B., Yu, J. Q., & Chen, Z. (2013). NBR1-mediated selective autophagy targets insoluble ubiquitinated protein aggregates in plant stress responses. PLoS Genet, 9(1), e1003196. doi:10.1371/journal.pgen.1003196

