## Supplementary figures for "Ambivalent roles of nonsense-mediated mRNA decay in heat stress tolerance": _suppFIGs_NMD-HS_20261030.pdf

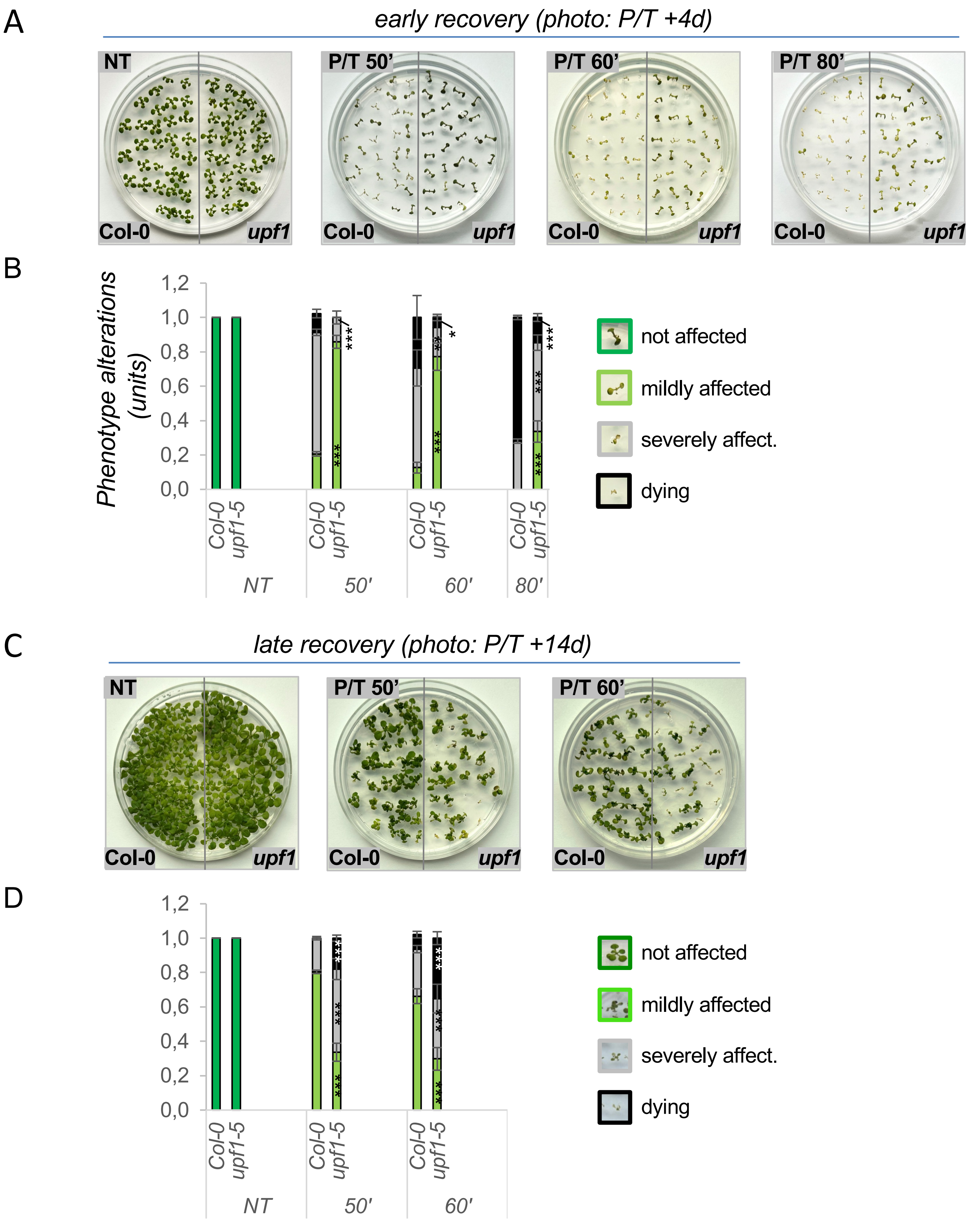


**Supplementary Figure S1: NMD pathway is needed for long term recovery following heat.** (A) phenotype of *upf1-5* mutant plants vs Col-0 following P/T treatment recovery 4 days (P/T+4d, early phenotype); triggering heat periods were 50, 60 or 80 minutes as shown; (B) quantification of early phenotype shown in *panel A*; (C) phenotype of *upf1-5* mutant plants following P/T treatment and 14 days recovery (P/T+14d, late phenotype); (D) quantification of late phenotype shown in *panel C*; triggering heat periods were 50 and 60 minutes as shown. Data are mean ± SD of three independent experiments, asterisks indicate significant differences to Col-0 (*$p < 0.05$; **$p < 0.01$; ***$p < 0.001$, unpaired two-sided t-test).

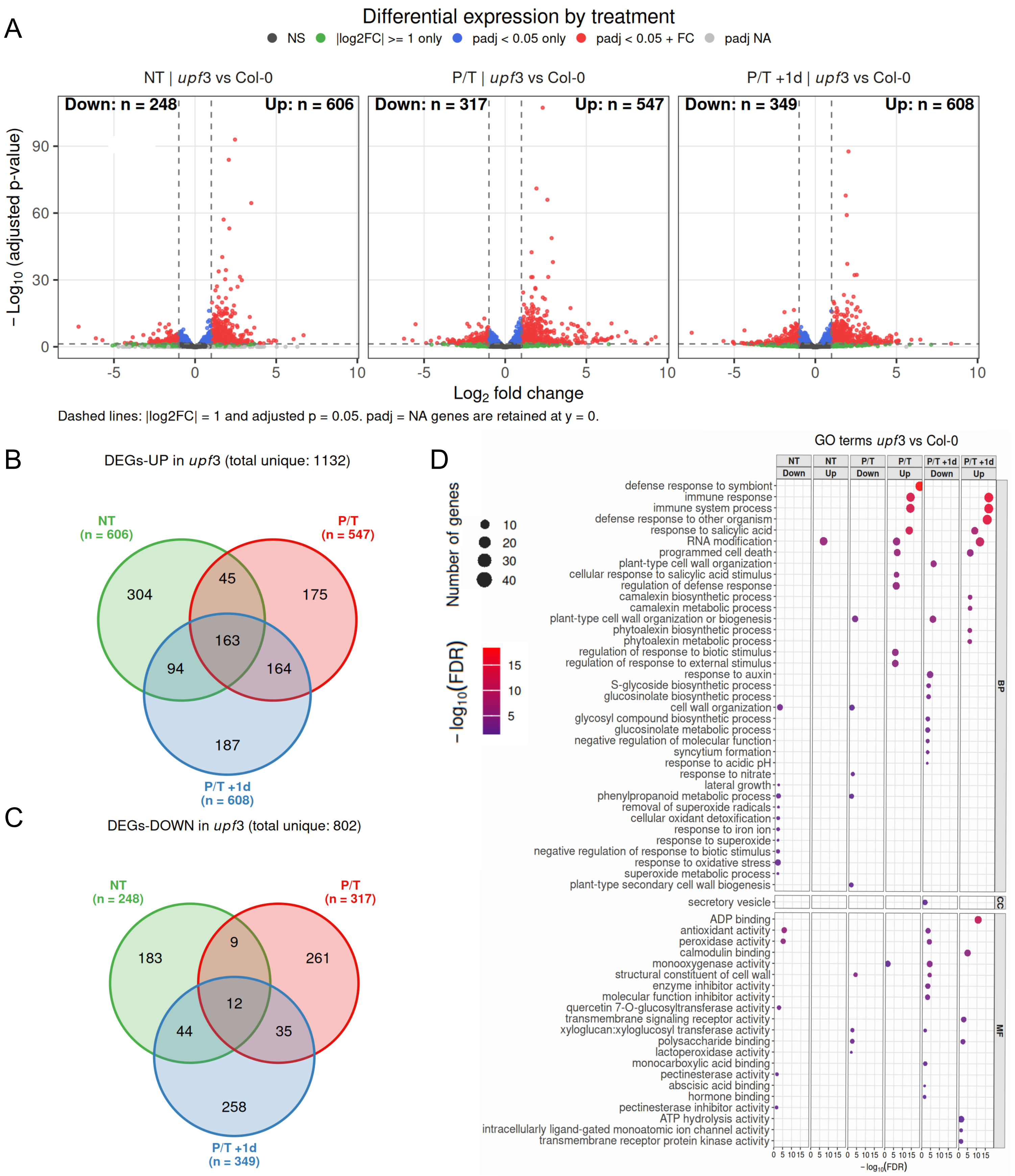


**Supplementary figure S2: RNA transcriptome changes of Col-0 and NMD mutant *upf3* plants in response to and following heat stress.** (A) Volcano plots of *upf3-1* vs. Col-0 wild-type differential expression at NT (*left*), P/T (*middle*), and 1 day recovery, P/T+1d (*right*); differentially expressed up- or down-regulated gene numbers ($p<0.05$, $\log_2 FC > 1$) are shown above each panel, with total transcripts tested (n=20127). (B) Venn analysis of differentially upregulated or (C) differentially downregulated genes in *upf3* vs Col-0 plants*,* gene numbers are indicated for each; (D) GO term enrichment across genotypes and conditions for genes down- or up-regulated in *upf3* vs. Col-0 at each condition (NT, P/T, P/T+1d), split by GO category, biological process (BP), cellular component (CC), and molecular function (MF) labels on right strips, GO terms are listed on the left, dot color indicates $-\log_{10}$(FDR) and dot size indicates the number of genes, as shown in the legend at bottom.

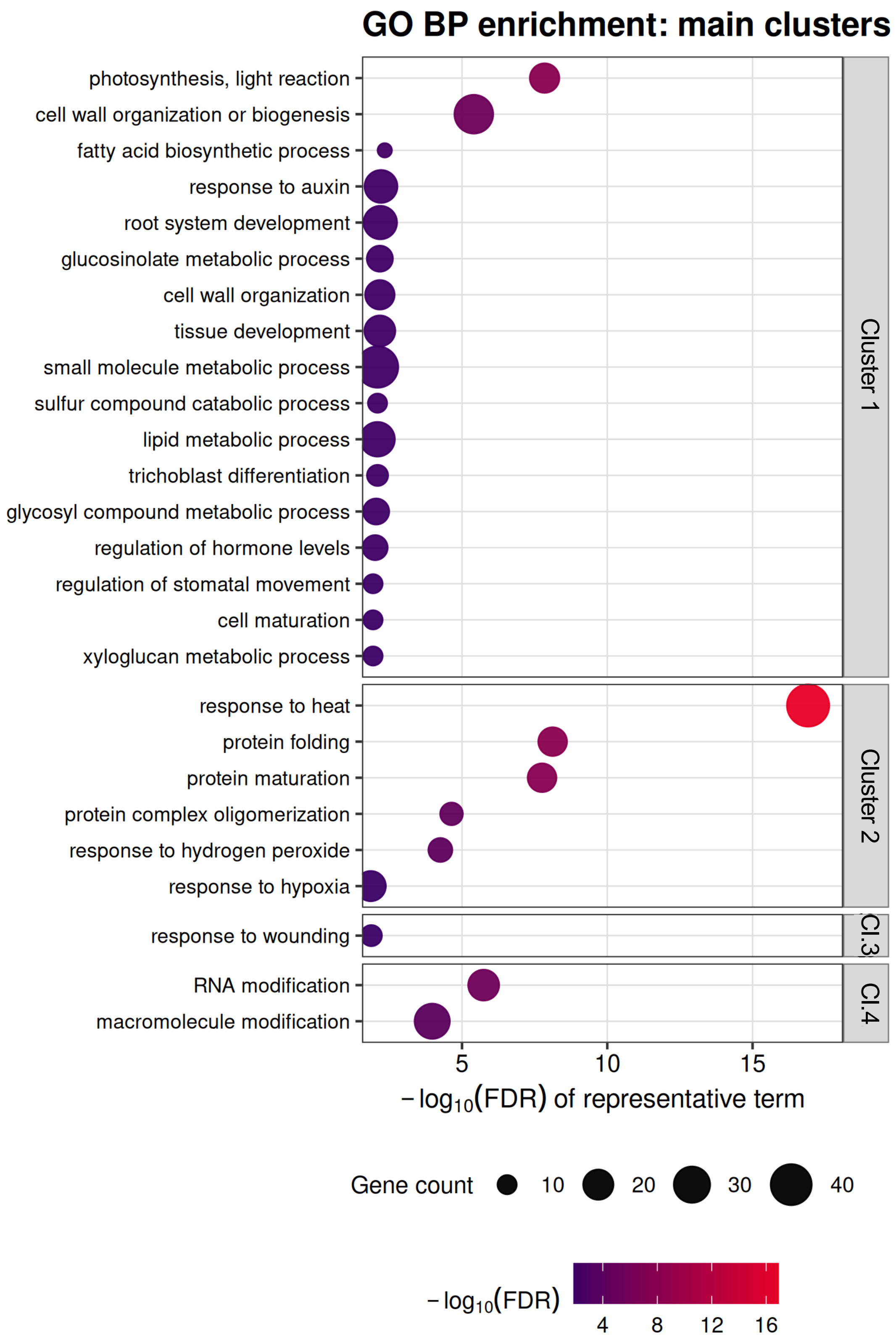


**Suplementary Figure S3. Gene Ontology (Biological Process) enrichment for Clusters 1–4**, unsupervised k-means trajectory clusters (k=6, *for cluster 5 see Fig 4C*, *subcluster 6 did not produced GO terms*). GO enrichment was computed with clusterProfiler (hypergeometric test, Benjamini-Hochberg-adjusted $p<0.05$); non-redundant terms were obtained by semantic-similarity reduction (rrvgo, threshold 0.7); dot size show gene count, color codes show significance, legend on the right side.

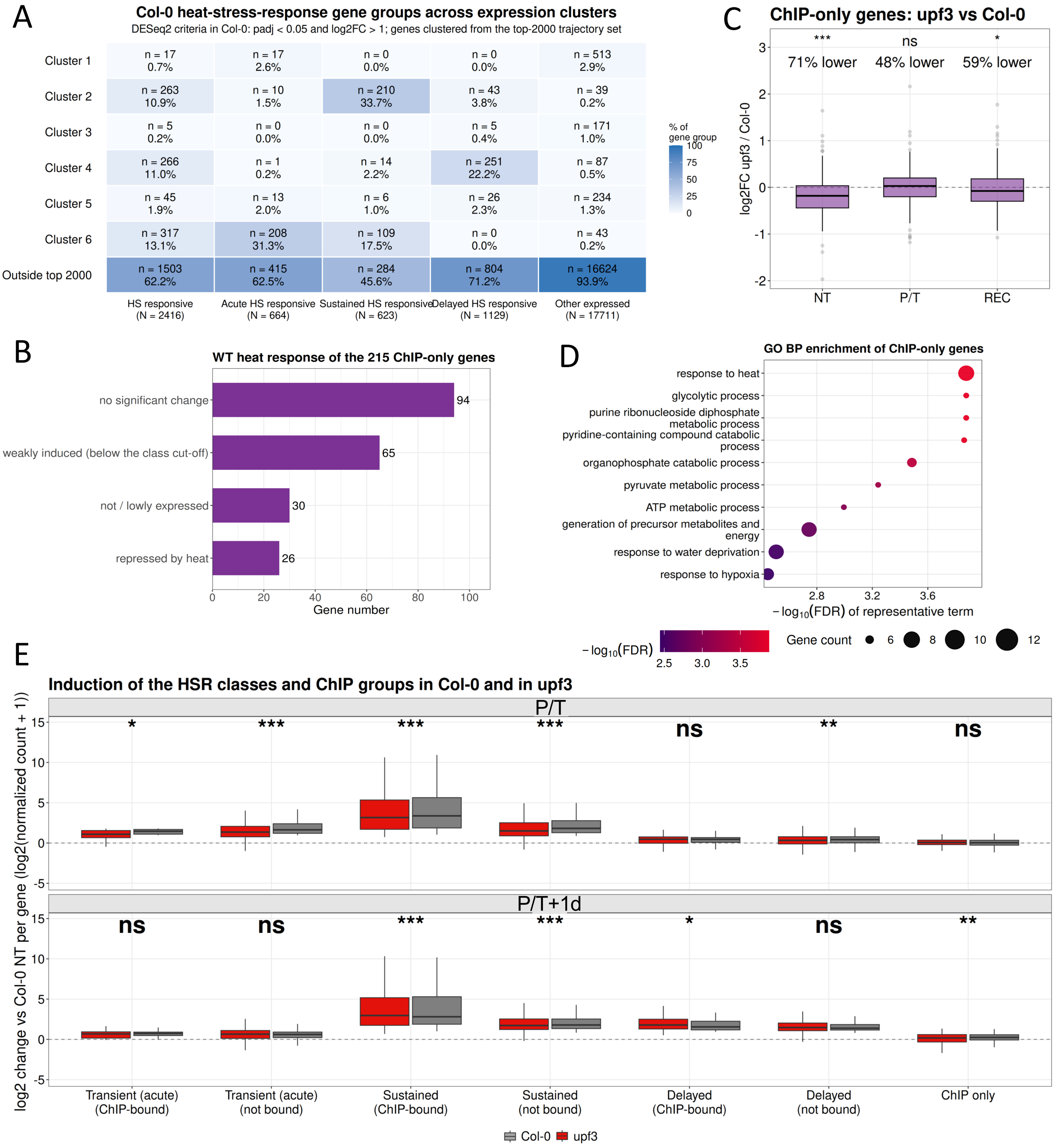


**Supplementary Figure S4. HSR gene class changes in *upf3*.** (A) Distribution of the Col-0 heat-stress-responsive (HSR) gene classes across the six expression clusters of the top-2000 trajectory set (padj < 0.05, $log_2FC > 1$): Each cell gives the number of genes and the percentage of that gene group. Genes not in the top-2000 set are shown in the bottom row. Colour shows the percentage of the gene group (*right*). (B) Col-0 HS changes of the ChIP-only set (genes bound by HsfA2/HsfA3 in the ChIP data but not part of the HSR classes). Bars show the number of genes grouped in different classes. (C) $log_2$ fold change of *upf3* relative to Col-0 for the ChIP-only genes at NT, P/T and REC. The percentages give the share of genes with lower expression in *upf3* than in Col-0; significance *p < 0.05, ***p < 0.001; ns, not significant; (D) GO BP enrichment of the ChIP-only genes, legend at *right*; (E) Induction of the HSR classes and ChIP groups in Col-0 (grey) and *upf3* (red) at P/T (*top*) and P/T+1d (*bottom*); HSR classes are split into ChIP-bound and not bound stes, the ChIP-only genes are shown separately. Significances of *upf3* vs Col-0 within groups *p < 0.05, **p < 0.01, ***p < 0.001; ns, not significant.

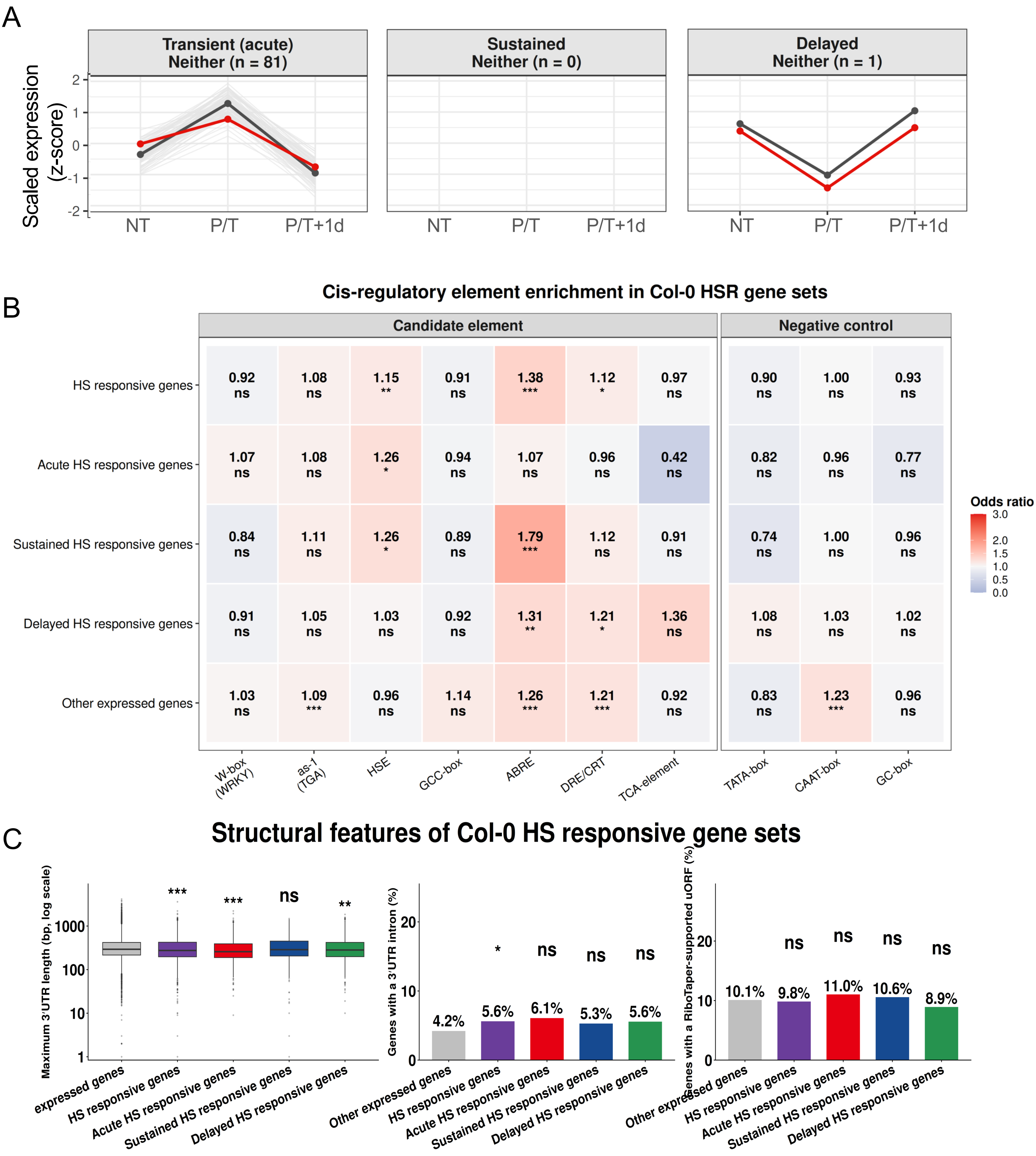


**Supplementary Figure S5. *Cis*-regulatory promoter elements and NMD structural features of the HSR genes.** (A) Expression trajectories at NT, P/T and P/T+1d of the HSR genes that are in neither of the groups defined in the figure, split by HSR class *control for main Fig 5C panels*. Grey lines are individual genes, and the thick lines are the class means in Col-0 (dark grey) and *upf3* (red). Values are scaled expression; (B) Enrichment of candidate *cis*-regulatory elements (W-box/WRKY, as-1/TGA, HSE, GCC-box, ABRE, DRE/CRT, TCA element) and negative-control motifs (TATA-box, CAAT-box, GC-box) in the HSR gene sets. Each cell gives the odds ratio, with colour scale from depletion (blue, <1) to enrichment (red, >1), and the significance of the enrichment (*p < 0.05, **p < 0.01, ***p < 0.001; ns, not significant); (C) Structural features of the HSR gene sets: 3′UTR length (bp, log scale), percentage of genes with a 3′UTR intron, and percentage of genes with a RiboTaper-supported uORF. Significance vs the other expressed genes (*p < 0.05, **p < 0.01, ***p < 0.001; ns, not significant);

A

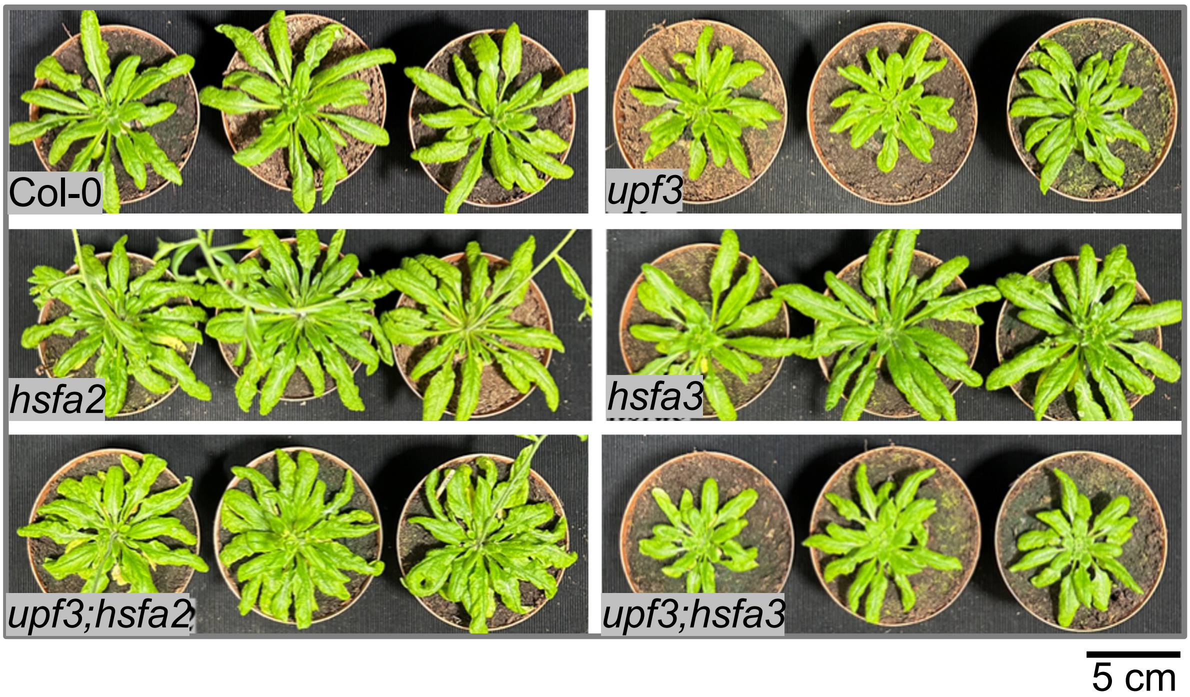


B

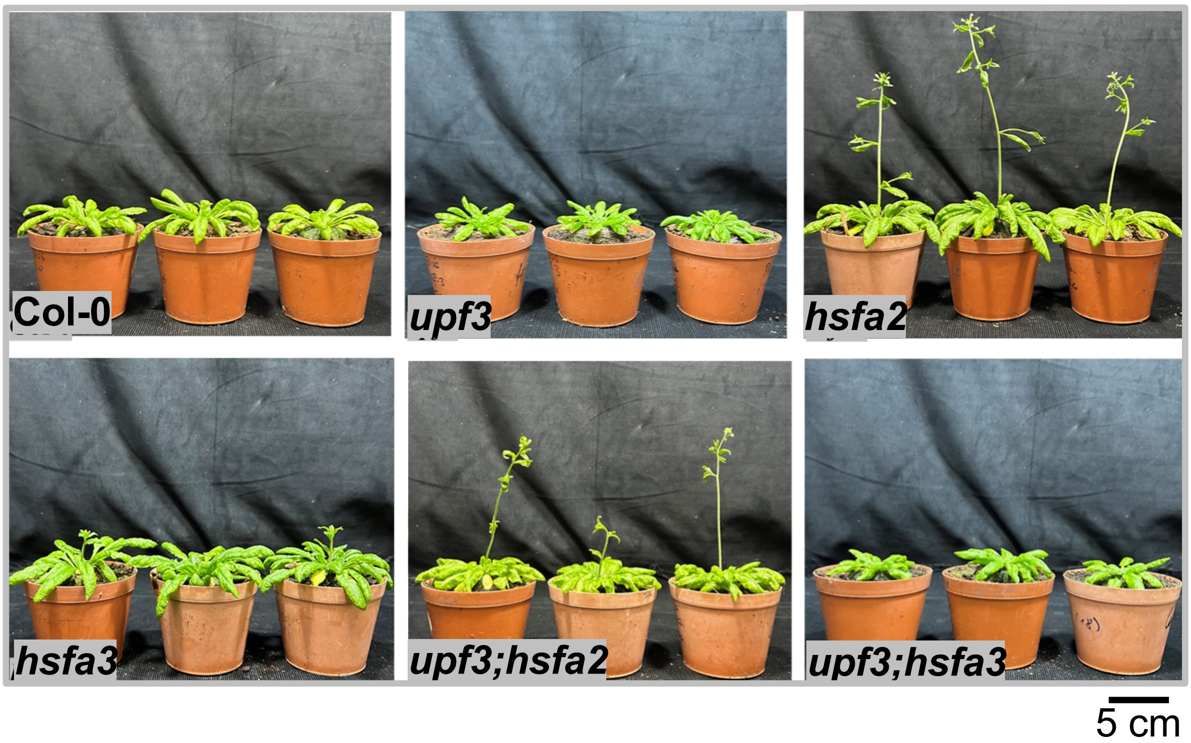


**Supplementary Figure S6. Developmental phenotypes of *upf3*, *hsfa2* and *hsfa3* single mutants and of *upf3;hsfa2* and *upf3;hsfa3* double mutants.** Col-0 (wild type) and the indicated mutants were grown under long day (16h light /8h dark) conditions at 21 °C without heat treatment and photographed at 30 days after sowing. (A) Top view of rosettes and (B) side view of plants; three plants per genotype are shown; *hsfa2* and *upf3;hsfa2* plants had started bolting at the time of imaging, whereas the other genotypes had not. Scale bar: 5 cm.

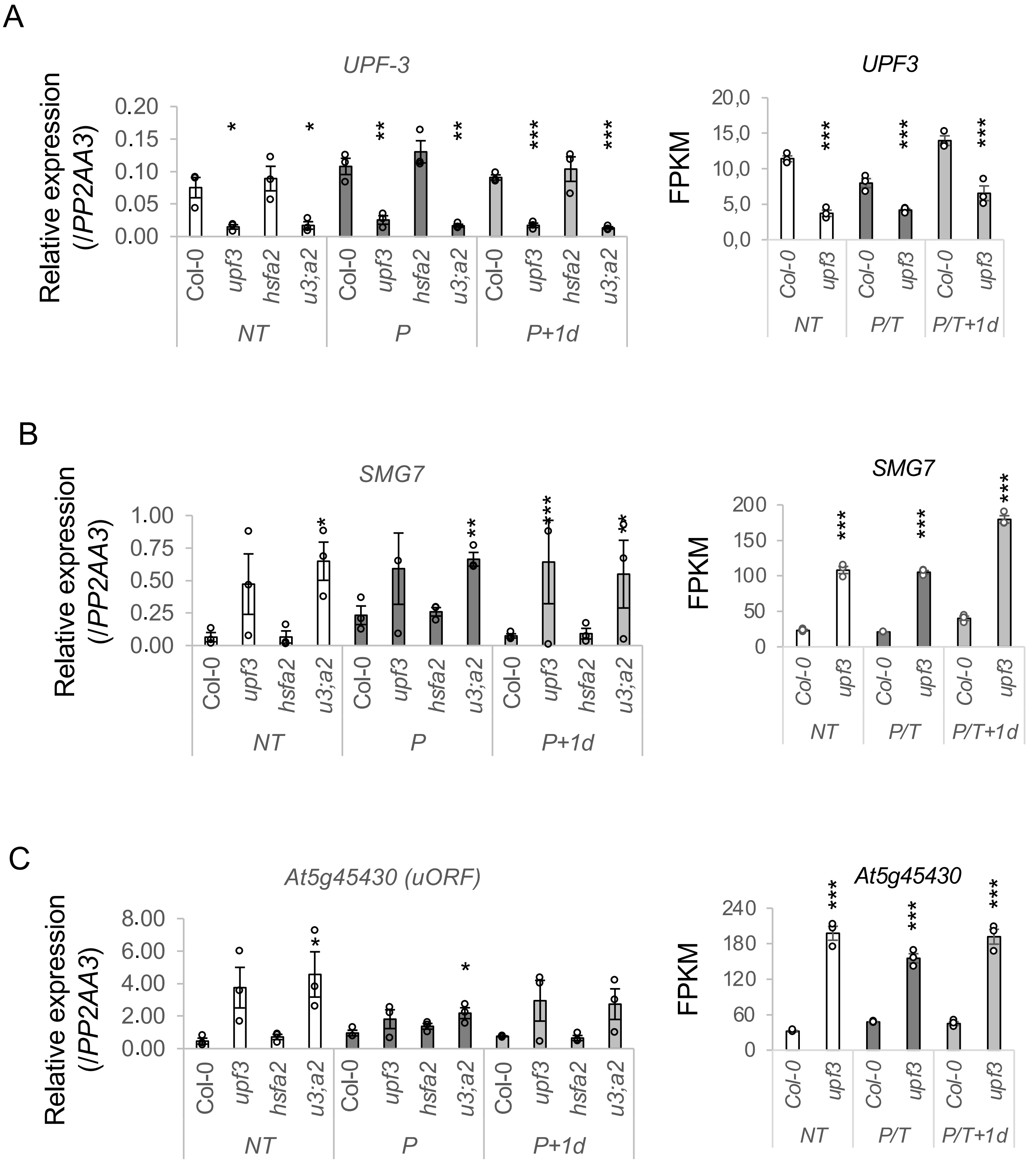


**Supplementary Figure S7. NMD remains active during and after heat treatment.** Left: RT-qPCR analysis of (A) *SMG7*, an NMD factor whose transcript is itself an NMD target, (B) *At5g45430*, an NMD target containing an upstream open reading frame (uORF), and (C) *UPF3* in Col-0, *upf3*, *hsfa2* and *upf3;hsfa2* (*u3;a2*) plants that were non-treated (NT), primed and triggered (P), or sampled one day later (P+1d); treatment regime as in Figure 1A. Expression is relative to *PP2AA3*; bars show mean ± SD of at least three biological replicates, and dots show individual values. Right: FPKM values of the same genes from the RNA-seq of Col-0 and *upf3* (NT, P/T and P/T+1d; mean ± SD of three biological replicates). Asterisks indicate significant differences between each mutant and Col-0 within the same condition (two-tailed, unpaired Student's t-test; $^*P < 0.05$, $^{**}P < 0.01$, $^{***}P < 0.001$). *UPF3* (C) is shown as a control for the *upf3* mutation.

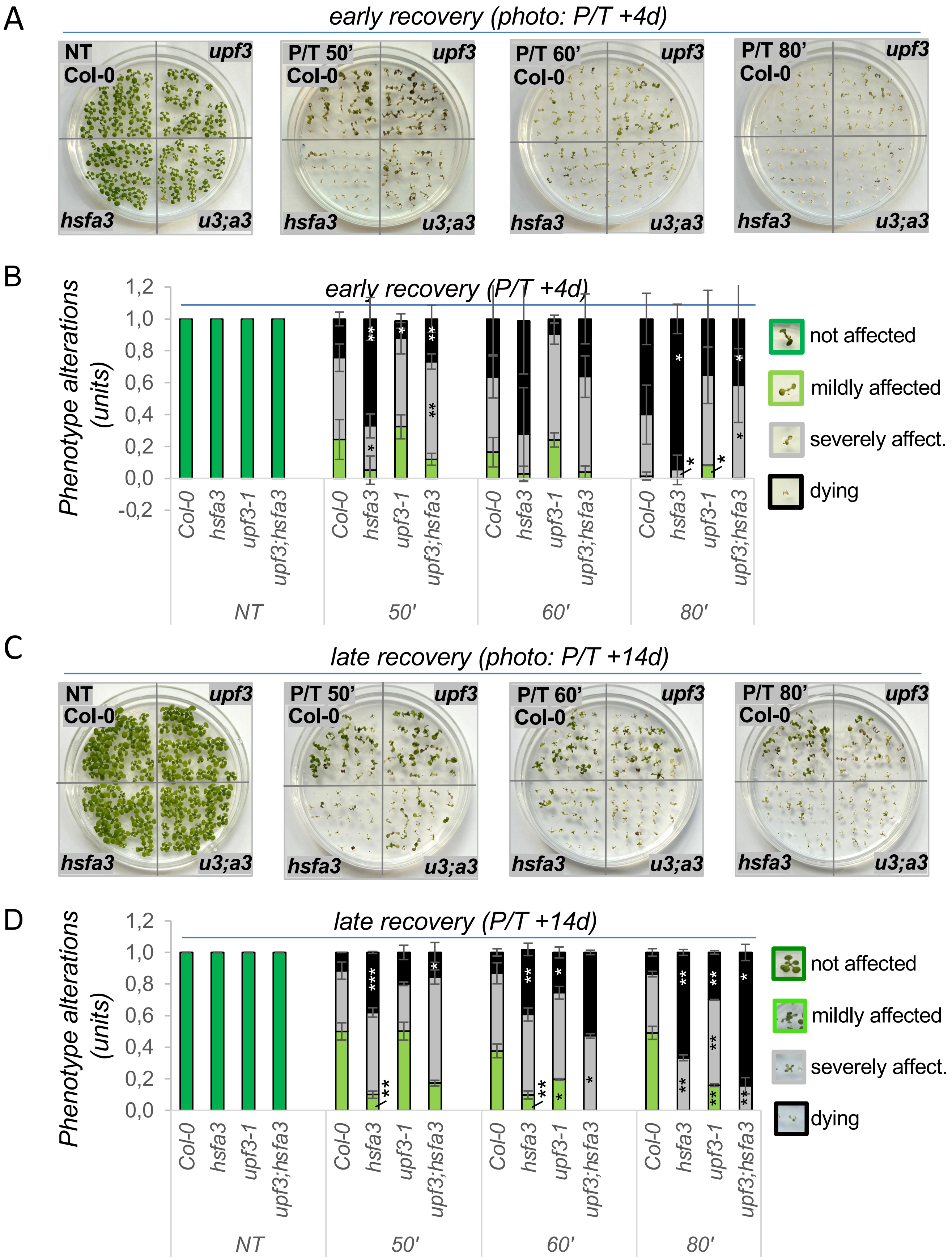


**Supplementary Figure S8: NMD's early impact on heat-stress recovery is independent of the HsfA2 pathway.** (A) Heat-stress phenotypes of wild-type (Col-0), *upf3*, *hsfa2*, and *upf3;hsfa2* (*u3;a2*) mutants on day 4 post-treatment (P/T+4d, early recovery); (B) Quantification of early-recovery phenotypes shown in *panel A*; (C) phenotypes of the same genotypes on day 14 post-treatment (P/T+14d, late recovery); (D) quantification of late-recovery phenotypes shown in *panel B*; data are mean ± SD of three independent experiments, and asterisks indicate significant differences of single mutants vs. Col-0 or of the double mutant vs. *hsfa2* (*$p < 0.05$; **$p < 0.01$; ***$p < 0.001$, unpaired two-sided t-test).

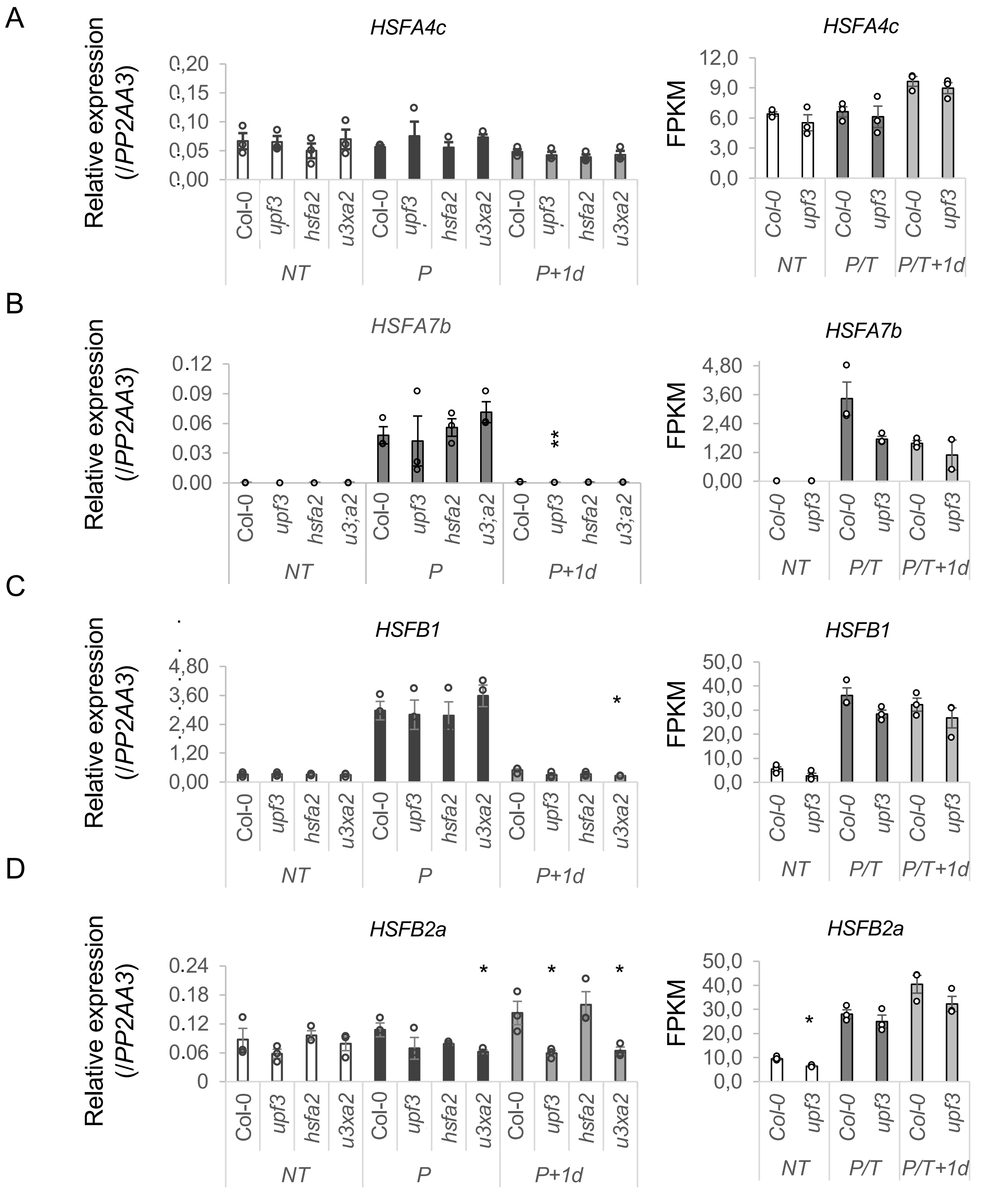


**Supplementary Figure S9. Expression of other heat-stress-responsive HSF genes is largely unaffected in *upf3*.** Left: RT-qPCR analysis of (A) *HsfA4c*, (B) *HsfA7b*, (C) *HsfB1* and (D) *HsfB2a* in Col-0, *upf3*, *hsfa2* and *upf3;hsfa2* (*u3;a2*) plants that were non-treated (NT), primed and triggered (P), or sampled one day later (P+1d); treatment regime as in Figure 1A. Expression is relative to *PP2AA3*; bars show mean ± SD of at least three biological replicates, and dots show individual values. Right: FPKM values of the same genes from the RNA-seq of Col-0 and *upf3* (NT, P/T and P/T+1d; mean ± SD of three biological replicates). Asterisks indicate significant differences between each mutant and Col-0 within the same condition (two-tailed, unpaired Student's t-test; $^*P < 0.05$, $^{**}P < 0.01$, $^{***}P < 0.001$); bars without asterisks are not significantly different.

A

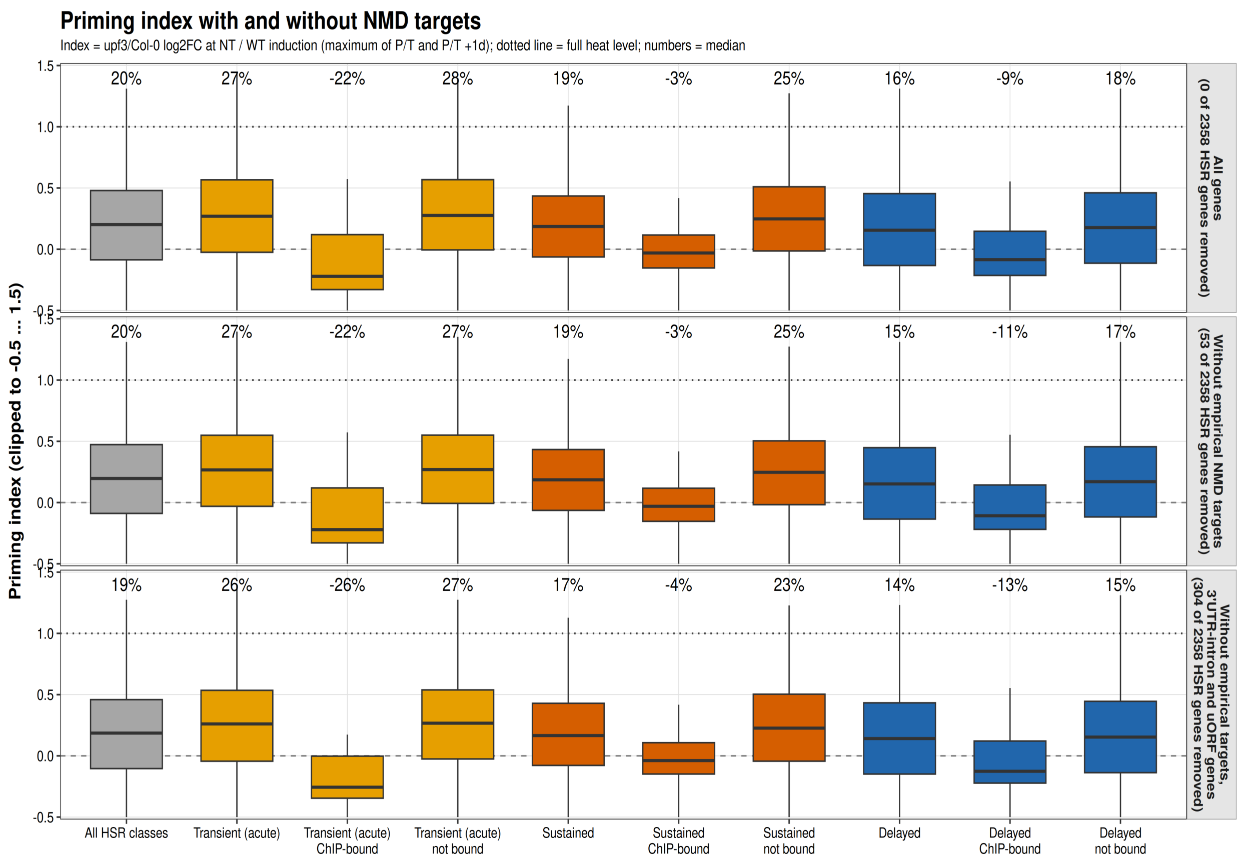


B

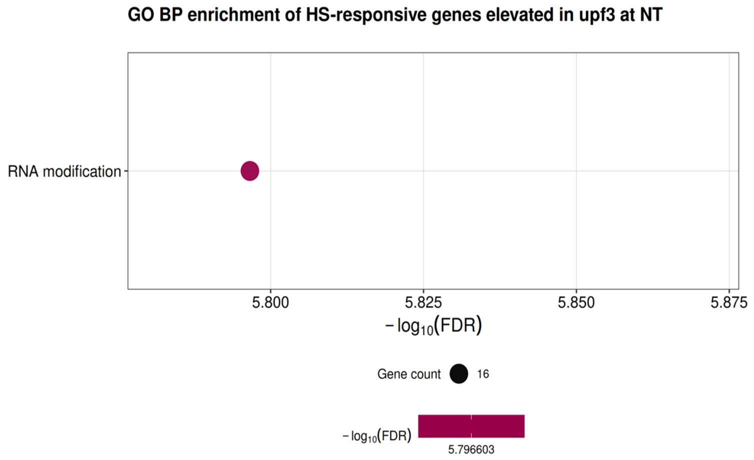


**Supplementary Figure S10. Analysis of NT priming startus in *upf3:*** (A) Priming index of the HSR genes at NT, with (*top*) and without NMD targets (*middle* and *bottom*). Priming Index is the *upf3*/Col-0 log2 fold change at NT divided by the maximum Col-0 induction (P/T or REC) of the same gene (PI = 0 means no difference from Col-0; PI = 1 (dotted line) means *upf3* at NT is at the full heat-induced level. Values are clipped to −0.5 to 1.5, and the number above each box is the group median. Gene groups are shown below; ChIPdata from Kappel et al. 2023. Rows show all HSR genes (n = 1,680), genes without empirical NMD targets (49 of 1,680 removed), and genes without empirical NMD targets and without a 3′UTR intron or uORF (277 of 1,680 removed); (B) GO biological process enrichment of the HSR-class genes that are elevated in *upf3* at NT, plotted by −log10(FDR), legend on the right.
